# Comprehensive analysis of αβT-cell receptor repertoires reveals signatures of thymic selection

**DOI:** 10.1101/2025.02.09.635277

**Authors:** Daniil V. Luppov, Elizaveta K. Vlasova, Dmitry M. Chudakov, Mikhail Shugay

**Affiliations:** Institute of Translational Medicine, Pirogov Russian National Research Medical University, Moscow, Russia; Institute of Personalized Oncology, I.M. Sechenov First Moscow State Medical University, Moscow, Russia; Department of Biological and Medical Physics, Moscow Institute of Physics and Technology, Dolgoprudny, Russia; ITMO University, Saint-Petersburg, Russia; Center of Molecular Medicine, Central European Institute of Technology (CEITEC), Masaryk University, Brno, Czechia; Department of Genomics of Adaptive Immunity Immunity, Shemyakin-Ovchinnikov Institute of Bioorganic Chemistry, Moscow, Russia; Center for Molecular and Cellular Biology, Moscow, Russia; Abu Dhabi Stem Cells Center, Abu Dhabi, United Arab Emirates

## Abstract

Thymic selection is a multi-stage process that establishes a T-cell immunity that is efficient in fighting diverse foreign pathogens, while not being self-reactive. During the process, T-cell receptor (TCR)α and β chains are rearranged to form a highly diverse set of heterodimers that are selected based on their affinity to peptides presented by major histocompatibility molecules (MHC) in the thymus. Here we employ high-throughput TCR sequencing data, theoretical model of TCR rearrangement and dedicated statistical methods to infer how the selection process affects human TCR repertoire on different scales. On a global scale, our results indicate differences in V(D)J gene usage, complementarity determining region 3 (CDR3) amino acid composition, CDR3 physicochemical properties, k-mer composition and differences in loop structure induced by the selection process. On a local scale, we were able to determine enriched TCR motifs and “holes” in repertoire induced by positive and negative selection and characterize their features. Finally, we demonstrated how TCR sequence composition affects lineage commitment via thymic selection and highlighted the effect of individual MHC haplotype. Our results can aid in identification of potentially self-reactive TCRs in donor repertoires in autoimmunity and immunotherapy studies.

**Graphical abstract (Figure 0):** **Study overview. A.** Datasets used in the study and the way TCR sequences were analyzed. **B.** Main directions in which thymic selection shapes TCR repertoire. Selection results in TCR-CDR3 losing positively charged and large amino-acids while increasing its flexibility. Unlike CDR3 of the TCRα chain, CDR3β hydrophobicity is also increased by selection. CDR3s carrying Cysteines and glycosylation sites are unlikely to pass the selection. In contrast, CDR3 carrying poly-Glycine regions are more likely to be selected for both chains. T-cells committed to CD8+ lineage were more likely to feature bulged CDR3s compared to CD4+. **C.** CDR3s enriched after thymic selection guide lineage commitment according to single-cell RNA sequencing data analysis, e.g. the MAIT cells and CD8+ phenotypes. **D.** Enrichment and depletion of certain TCR motifs pinpoint fine-structure of post-selection repertoires, as inferred by sequence neighborhood analysis. **E.** Comparative analysis of enriched and depleted TCR clusters in monozygotic twins revealed that the selection process is shaped by HLA haplotype.

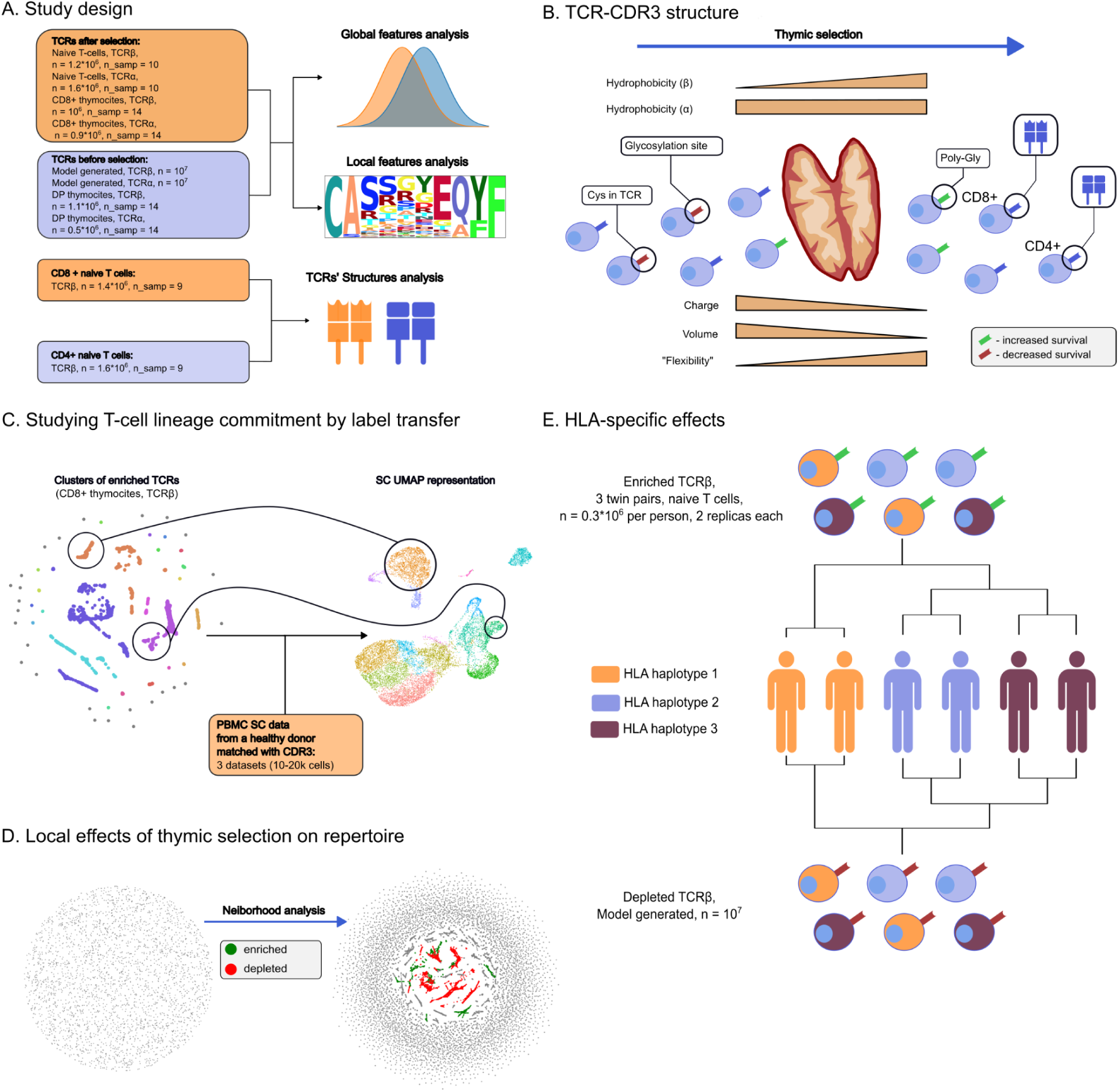

## Introduction

Lymphoid progenitors migrate from bone marrow to thymus where they develop their T-cell receptor (TCR, a heterodimer of α and β chains) critical for antigen recognition via a process called Variable-(Diversity)-Joining gene rearrangement and undergo screening and selection in order to become mature T-cells. The aim of this selection is to eliminate thymocytes without the ability to recognise antigens presented by major histocompatibility complexes (MHCs) on other cells (positive selection) and thymocytes whose affinity to the complexes is too strong and may cause self-recognition (negative selection). Thus, TCR sequence is the main factor of passing thymic selection and it affects lineage commitment and the overall future of a T-cell (Duan and Mukherjee, 2016).

The process of V(D)J gene rearrangement involves two main steps: 1) three (V, D, and J for TCR β chain) or two (V and J for TCR α chain) gene alleles from corresponding loci are selected and recombined together, 2) bases are randomly deleted at gene ends and non-template (N) nucleotides are added to the junction sites between genes to increase sequence diversity. The abundance of different genes to be combined together along with a random nucleotide inclusion/deletion results in a great variety of possible TCRs, estimated to be ∼10^19^ (Dupic et al., 2019), which allows TCRs to recognise enormous diversity of antigens potentially presented by MHCs, and thus to reliably recognize a multitude of antigens from each specific pathogen with a diverse set of receptor variants. This diversity is also orders of magnitude greater than the total number of T-cells in the human body (∼10^11^, (Farber et al., 2014).

Before a T-cell is released into bloodstream it should pass the thymic selection in order to ensure that it is both functional in terms of antigen recognition and won’t mistake healthy cells for infected or malignant ones (Duan and Mukherjee, 2016). Thymic selection is a multi-staged process (Robert et al., 2021) in which CD4-CD8- double-negative (DN) T-cells rearrange β chain, validate its functionality in the complex with pre-TCRα, then rearrange α chain and acquire CD4+CD8+ double-positive (DP) phenotype together with a mature TCR that is subject to positive and negative MHC-guided selection; these DP T-cells then differentiate to either CD4+ (helper or Treg) or CD8+ (killer) single-positive (SP) phenotype and emigrate from thymus.

The most variable region of TCR is the complementarity-determining region (CDR) 3 encoded by the V(D)J junction that is in direct contact with the peptide presented by MHC. Two other complementarity-determining regions - CDR1 and CDR2 are far less diverse and encoded by V gene, they are also responsible for peptide recognition and may engage the antigen binding groove of MHC (Bosselut, 2019). The high diversity of CDR3, the fact that it contains info on V and J genes, and its direct contact with an antigen is the reason why it usually serves as a proxy for “TCR sequence” in most recent studies (Rosati et al., 2017). In present study we mostly focus on CDR3 sequences when describing the thymic selection process from the TCR repertoire perspective.

Most modern TCR repertoire studies are based on high-throughput sequencing technologies capable of reading millions of TCR sequences from bulk samples and single cells, a methodology collectively known as Rep-Seq (Benichou et al., 2012) or AIRR-seq (Rubelt et al., 2017). To this date, AIRR-seq has been applied to numerous biological samples, coming from different donors, tissues and conditions, including thymocytes, naive T-cells, sorted T-cell populations and bulk peripheral blood mononuclear cells (PBMCs). Since there are quite few previously published TCR repertoires from human pre-selection thymocytes we and others employed a theoretical model of V(D)J rearrangement to simulate pre-selection TCR sequences (Sethna et al., 2019). This model was developed based on recombination events probabilities inferred from non-functional TCRs, which was not subjected to any functional selection and thus capable of representing DN repertoire before thymic selection. We also rely on the hypothesis that “singleton” T-cell clonotypes supported by a single mRNA molecule in a deep repertoire predominantly albeit not exclusively represent naive T cells in order to model post-selection naive TCR repertoire as released from the thymus (Britanova et al., 2014).

We note that as the majority of present AIRR-seq datasets contain TCRβ chain sequencing data, we focus our analysis on it and additionally analyze TCRα chain to highlight specific features where possible. One of the excuses for focusing on the TCRβ chain only is that, as mentioned previously, it rearranges first and many suggest that it is detrimental for T-cell fate (Rothenberg, 2023). The notable exception here are single-cell datasets used in present study that have information on TCR chain pairing, however, those are quite limited in the quantity of T-cells sequenced.

Our general approach in present study was to compare TCR repertoires before and after selection using model data and conventional PBMC AIRR-seq data and further validated our findings using sorted DP and SP thymocytes (Quiniou et al., 2023). We explore both local and global repertoire structure of pre- and post-selection TCRs by analysing sequence features that can influence the selection process on various levels: V/J gene usage, amino acid composition and physicochemical properties of primary CDR3 sequences, CDR3 k-mer profiles of repertoires and prominent sequence motifs. In order to refine comparative analysis and remove noise originating from intrinsic randomness of V(D)J rearrangement we utilized a TCR sequence cluster enrichment strategy based on the TCRNET method, allowing us to detect selection motifs in whole-body TCR repertoires with intrinsically complex structure (Pogorelyy and Shugay, 2019).

Some of these TCR features were reviewed in the context of thymic selection in previous studies. For example, study by Lu et al. (Lu et al., 2019) revealed notable differences in amino acid usage before and after thymic selection in MHC-restricted mice repertoires, such as elimination of hydrophobic, positively charged amino acids and Cysteines. In concurrence with the latter study Stadinski et al. (Stadinski et al., 2016) demonstrated that TCRs with hydrophobic residues in particular positions are prone to cross- and self-reactivity, thus lacking chances to survive thymic selection. Other studies revealed local repertoire features differences between T cell repertoires in thymus, distinct lymph nodes, and spleen (Feng et al., 2015; Nakonechnaya et al., 2024). These features were also revisited in Isacchini et al. using repertoire modeling, yet the study arrives at the conclusion that selection features resemble themselves on local and global scales, claiming that there are no forbidden TCR sequences and selection motifs. We review the aforementioned features using our framework and arrive at similar conclusions, yet we extend previously published findings by showing locally enriched and depleted TCR motifs that can be both public and donor/HLA-specific.

Linkage between lineage commitment and TCR was extensively studied, revealing distinct CDR3 features between CD4 and CD8 (Camaglia et al., 2023), CXCR3+ and CXCR3- naive CD8 (De Simone et al., 2019) and helper T cell subsets (Kasatskaya et al., n.d.). Here we explore it in more detail via single-cell data, showing that selection motifs are linked to certain phenotypes. We demonstrate that TCRs having characteristic features inferred for CD8+ thymocytes are mostly carried by CD8+ T-cells in donor samples. Moreover, we show that the structure of the CDR3 loop is different for CD4- and CD8-related TCR motifs. Additionally, it is known that non-canonical T-cells are present in the thymus and show specific TCR features. Among them there are Mucosal-associated invariant T cells (MAIT) cells (Godfrey et al., 2019) which carried semi-invariant TCR. In our analysis we managed to trace some trivial cases of lineage commitment and link them to TCR CDR3 structure as well.

We also considered the role of donor HLA haplotypes in selection and demonstrated how allele-specific differences result in global repertoire features. Such effects had been already reported for mice CD4+ repertoires (Logunova et al., 2020). It was also shown that MHC context generally shapes the T-cells repertoire (Ishigaki et al., 2022). Similar to our study approach, which involves twins’ TCR repertoire, was used by Tanno et al. to reveal genetic factors, in particular MHC alleles, impact on TCR repertoire (H et al., 2020).

## Materials and Methods

### Post-selection T-cell repertoires

We used previously published peripheral blood mononuclear cell (PBMC) TCR repertoire sequencing data for both TCRα (Heikkila et al. 2021) and TCRβ (Emerson et al., 2017) chains. For TCRβ, a sample of 10 CMV donor repertoires selected *ad hoc* were chosen from the HIP cohort of Emerson *et al*. dataset for TCRβ analysis (sample IDs in dataset from 1 to 10). Only clonotypes which are supported by single read (singletons) were used in our analysis: singletons serve as a good proxy for naive T-cells since as their fraction is almost equal to the naive population size estimated by flow cytometry and their sampling behavior is in line with what is expected from rare naive cells (Britanova et al., 2014).

### Sorted repertoires

Repertoires of sorted CD4+ and CD8+ naive (post-selection) T-cells sequencing were taken from Qi *et al*., this data is available for TCRβ chain sequencing only. TCR repertoire for double positive (DP) and CD8+ (single-positive, SP) thymocytes (pre-selection) were taken from a recent Quiniou et al. study. Note that for these datasets no read count information was used (i.e. all clonotypes were assumed to be singletons) in order to avoid potential amplification biases and make it compatible with other datasets.

### HLA matched and mismatched repertoires (twin studies)

A dataset containing PBMC TCRβ repertoire sequencing for three pairs of twins was taken from Pogorelyy *et al*. (Pogorelyy et al., 2018). Pre-vaccination and day 0 repertoires sampled prior to treatment were used as biological replicates for each twin, only singletons were included in the analysis Additional dataset containing PBMC TCRα and β repertoires from three pairs of twins was taken from Zvyagin *et al*. (Zvyagin et al., 2014). Sorted naive CD4+ T-cells TCRβ repertoires from two pairs of twins were also obtained from Kasatskaya *et al*. (Kasatskaya et al., n.d.). Summary statistics for datasets are reported in Supplementary Tables 1, 3 and 4.

### Simulating pre-selection V(D)J rearrangements

Pre-selection TCRα and β repertoires were simulated based on a theoretical probabilistic model of the V(D)J rearrangement process using OLGA software (v1.2.4) as described previously (Sethna et al., 2019). The software was executed with default runtime parameters and model probabilities, random seed was set to 100 and a sample of 10^7^ random rearrangements was generated for each of TCR chains.

### Single-cell data analysis

We utilized a standard 10X Genomics datasets containing 20,000 PBMCs of a healthy donor (https://www.10xgenomics.com/resources/datasets/20-k-human-pbm-cs-5-ht-v-2-0-2-high-6-1-0), however, our main findings can be reproduced using other 10X PBMC datasets containing both TCR repertoire and RNA sequencing data.

Single-cell RNA sequencing data analysis was performed using Seurat R package (version 4.9.9.9058) (Hao et al., 2021). Data preparation, dimensionality reduction and UMAP visualization were carried out according to commonly used guidelines for 10X data: (https://satijalab.org/seurat/articles/pbmc3k_tutorial.html). Differential expression analysis for T-cells carrying TCR sequences of interest was performed using MAST algorithm (Finak et al., 2015) implemented in Seurat package.

For additional Single Cell data analysis datasets totaling 178307 cells of 88 healthy patients were taken from Lindeboom et al work (Lindeboom et al., 2024). Cluster abundance on a particular cell type was tested by Fisher exact test.

### TCR amino acid sequence feature and motif analysis

Basic features of the TCR sequence such as V/J gene usage, single amino acid frequencies, k-mer (k=3) frequencies and physicochemical of CDR3 regions were carried out using in-house scripts as described previously (Shugay et al., 2015). Kidera factors (Kidera et al., 1985), charge and hydrophobicity were calculated using “peptides” python package (v0.3.2).

Detection of TCR CDR3 sequence motifs was performed using TCRNET algorithm implemented in VDJtools (v1.2.1) as described previously (Pogorelyy and Shugay, 2019). Briefly, this method defines TCR sequences of interest as those, that are placed in the more dense regions of CDR3 sequence similarity graph compared to a control (typically produced assuming V(D)J rearrangement model with no selection pressure) dataset: the number of 1-hamming distance neighbors is compared to the expected number of neighbors adjusted for sample and control sizes producing an enrichment score and a P-value based on Binomial approximation.

Note that in order to infer TCR sequence clusters that were depleted by negative thymic selection, we simply swapped “background” (control) and “foreground” (our sample of interest), i.e. we searched for TCRs enriched in pre-selection data compared to post-selection.

In order to produce a representative set of TCR clusters, we selected top 10,000 neighbor-enriched CDR3 sequences based on enrichment P-value. Motifs (CDR3 sequence logos) for selected clusters were visualized using logomaker package (v0.8). The top 5 largest clusters were subsequently analyzed. The number of clusters was selected *ad-hoc*. SoNNia (v0.2.3) model was additionally used to assess difference in amino acids’ occurrence probabilities in particular position in post- vs pre- selection repertoires (Isacchini et al., 2021).

### Comparative analysis of twins dataset

In order to identify positively and negatively selected CDR3 clusters in the twins dataset, we subsampled each twin sample to 306,553 CDR3s (size of the smallest repertoire) and pooled all the samples together. We used *ad hoc* thresholds to select significantly enriched (log2 fold change > 2 and -log10 p > 12 for sample pool compared to simulated sequences as control) and depleted (log2 fold change > 1 and -log10 p > 12 for simulated sequences compared to sample pool as control) clusters. Clusters containing more than 10 sequences were used in further analysis. Next, similarity between positively and negatively selected clusters was estimated by computing Jensen-Shannon divergence between cluster frequencies defined as the number of clonotypes from a given cluster present in a given sample. Jensen-Shannon divergences were calculated via scipy package (v1.10.1).

### Structure analysis

CDR3 loop structures for TCRs of interest were modeled using TCRmodel web tool (Yin et al., 2023) and processed with Pymol (version 2.3.0). Our in-house “mir” software package was used to annotate resulting PDB files (see (Karnaukhov et al., 2024)). As CDR3α is not known in most of our datasets, we used a generic CAGGSSNTGKLIF (TRAV27, TRAJ37) sequence that is most commonly observed variant in the Heikkila *et al*. dataset as a dummy TCRα sequence. TCR CDR3 backbone was visualized by applying PCA to Cα atom coordinates using sklearn package (v1.3.1).

### Code availability

All code used in this study is available at https://github.com/LuppovDaniil/Thymic_selection_notebooks (Python version 3.11.5., R version 4.1.2).

## Results

### Comparing pre- and post-selection TCR amino acid sequences

AIRR-seq data for DP T-cells sorted from thymus can be used to explore the initial space of rearranged TCRα and β sequences existing prior to positive and negative, similar to the work of Quiniou *et al*. (Quiniou et al., 2023). Alternatively, recent studies have shown that the structure of V(D)J rearrangement space can be accurately recaptured by a rather simple probabilistic model that allows sampling TCR sequences similar to those produced *in vivo* in both their amino acid composition and frequency (Sethna et al., 2019). Here we use both Quiniou *et al*. and model datasets as a pre-selection repertoire, but must note the former lacks data on TCRα while the latter doesn’t account for β selection in DN cells that occurs when pre-TCR is formed.

There are several ways to acquire the TCR repertoires of post-selection T-cells that have not yet undergone strong antigen exposure and obtained memory phenotype. One can either sort and sequence naive CD4+ and CD8+ T-cells as performed in Qi *et al*. (Qi et al., 2014), or use SP thymocytes as done in Quiniou *et al*. (Quiniou et al., 2023). Alternatively, one can select T-cell clonotypes detected only once (singletons) from unsorted PBMC AIRR-seq data, as they mostly represent naive T-cells (see Britanova et al., 2014). In the present study we used Qi *et al*. and Quiniou *et al*. datasets, and selected singletons from 10 samples chosen *ad hoc* from the Emerson *et al*. dataset, leading to approx. 10^6^ CDR3 variants.

Thus, in this study we took repertoire sampled from a recombination model (namely OLGA (Sethna et al., 2019)) as a repertoire before the selection and singletons repertoire from PBMC as a repertoire of naive T-cells after selection. Additionally, thymocytes DP and SP repertoires were taken in a parallel analysis (Graphical abstract 1A).

Individual amino acid frequency analysis in TCRβ CDR3 revealed substantial decrease in usage of some of them in post-selection repertoires compared to expected from V(D)J rearrangement model (Figure 1A) in agreement with previous observations (Elhanati et al., 2014). Three of them, Arginine, Histidine and Lysine are positively charged and physically large amino acids that may result in negative selection due to strong antigen binding or sterical difficulties in antigen recognition (Kosmrlj et al., 2008). Proline and Cysteine are negatively selected due to their impact on protein and loop structure: while the former can result in turns that can significantly alter CDR3 omega loop structure, the latter may form disulfide bridges with Cysteines in the Variable region and disrupt folding. These trends are generally reproduced in experimental thymocytes data (Supplementary Figure 1A).

**Figure 1.**
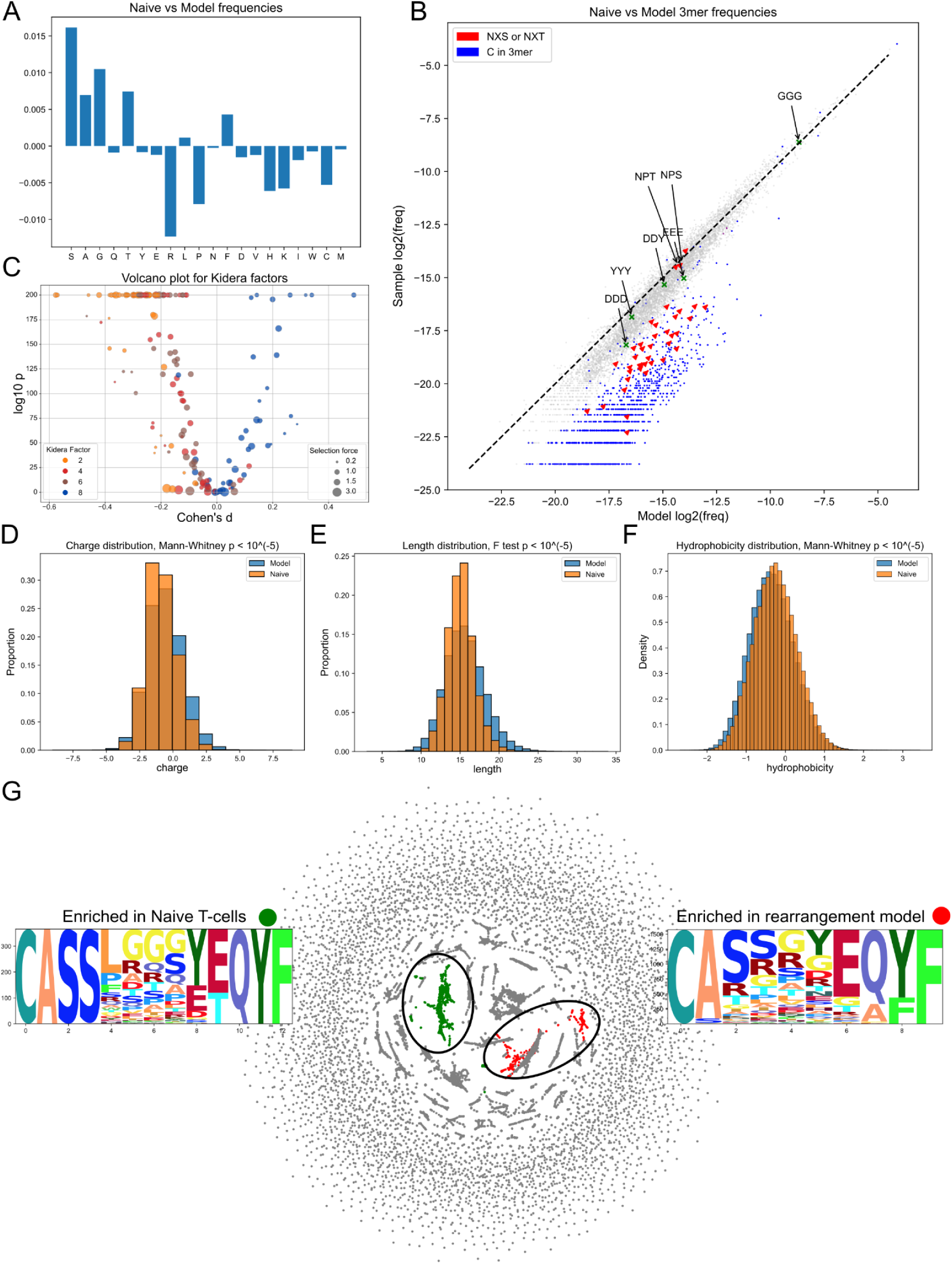
Model generated CDR3β and naive CDR3β repertoire (naive sample) comparison. **A.** Single nucleotide frequencies before and after selection comparison. **B.** 3-mers frequencies comparison. Glycosylation sites (red) and C-containing 3-mers (blue) are negatively selected. Dashed line represents the y = x relationship. **C.** Volcano plot for Kidera factors affected by selection. Each point represents a VJ pair. Negative shift in Kidera factors 2, 4 and 6 and positive shift in Kidera factor. The ratio of fractions of a particular VJ pair before and after selection were labeled as “selection force”. D. Negative shift for charge distribution after selection. **E.** “Winsorizing” of CDR3 length values post selection. **F.** Positive shift in hydrophobicity values after selection. **G.** Largest enriched and depleted clusters and their logos obtained by sequence neighborhood analysis.

3-mer frequencies analysis showed that 3-mers with Cysteine had less chance to survive selection (Fig. 1B) as we observed it on a single amino acids level (Fig. 1A). 3-mers containing sequences NX[S,T], where X can be any amino acid, are similarly less likely to survive the selection (Fig. 1B). Notably, such sequences represent sites of N-glycosylation (Marshall, 1972). Among all of the NX[S,T] sequences the top 3 least affected by selection 3-mers includes NPT and NPS. (Fig. 1B). This is in line with the observation that glycosylation does not occur at these sites (Sun et al., 2019). Glycosylation was previously reported in the context of B-cell receptor (BCR) and Follicular lymphoma (Zhu et al., 2002) but not in the context of TCRs. Additionally, we checked sulfonation sites motifs which were described in (Pospelova and Safonova, 2022) for antibodies, such as DDD, DDY, YYY and EEE. The tangible reduction effect was observed only for DDD motif. Both glycosylation sites and Cysteine linked effects were observed in real-world data as well as the lack of effect from putative sulfation sites (Supplementary Figure 1B).

In order to describe changes in physicochemical properties of TCRβ CDR3s we harnessed Kidera factors (Kidera et al., 1985). These factors represent the key physical properties of amino acids obtained by dimensionality reduction in a space of their physicochemical properties. We compared each VJ pair independently in order to lessen the bias caused by the choice of the V and J genes (Fig. 1C). Among Kidera factors negatively affected by selection were Kidera 2, which determines side chain size, Kidera 4, which is inversely associated with hydrophobicity, and Kidera 6, which determines partial specific volume. These results suggest that TCRβs with a physically smaller hydrophobic CDR3 have a greater chance of passing selection. The only Kidera that changed positively was Kidera 8 - occurrence in the α-helix structural region. Notably, the selection effect within a particular Kidera Factor was shared among all VJ pairs. Identical analysis of DP and SP thymocytes mirrored the above results, however, the effect for Kidera factor 6 was less pronounced (Supplementary Figure 1C).

General physical properties of TCRβ CDR3 were assessed without separation to VJ pairs. We observed that a charge of repertoire has decreased after selection (Fig. 1D). Such observation is expected since drop in frequencies of positively charged amino acids was also observed (Fig. 1A). The distribution of lengths demonstrated the effect of some sort of winsorizing - too short and too long sequences both have less chance to get through selection (Fig. 1E). Unlike charge, hydrophobicity markedly increased after selection (Fig. 1F). Except for length, which experienced shortening instead of winsorizing, the same effects were detected on real-world thymocytes data (Supplementary Figure 1D-F).

Next, we assessed functional clusters of TCRβ CDR3s enriched in naive cells compared to the modeled sample and vice versa, using VDJ Tools (Shugay et al., 2015). Largest enriched clusters in naive and model-generated samples are presented in Fig. 1G. Model-enriched clusters can be interpreted as sequences which tend not to pass the selection and naive sample-enriched clusters as ones which in contrast are likely to survive thymic selection.

The majority of the post-selection enriched clusters contain poly-Glycine sequences in the middle of a TCRβ CDR3 (Fig. 1G, Supplementary Figure 2) that is known to be one of the most flexible among the polypeptides chains (Bykov and Asher, 2010). K-mers analysis demonstrated that GGG 3-mer passed through the selection with unchanged frequency in accordance with the cluster analysis (Fig 1B, Supplementary Figure 1B). Clusters enriched in pre-selection samples carried Arginine and Proline, which may have a strong impact on the CDR3 loop structure. The consensus sequence CASS at the beginning of the CDR3 were often aberrant in these clusters (Fig 1G, Supplementary Figure 2). Notably, clusters enriched in DP and SP thymocytes mostly resembled clusters enriched in model-generated and naive repertoires respectively (Supplementary Figure 3). Latter highlights that for TCRβ our analysis is reproducible.

In order to obtain pre- and post- selection probabilities of amino acids occurrence in the particular position in CDR3 we utilized the SoNNia software (Isacchini et al., 2021). Generally, we observed the nearly identical to enriched clusters analysis trends for both of the repertoire-pairs being analyzed (Supplementary Figure 4A, Supplementary Figure 4C).

Gene usage analysis revealed a vague picture of little or no preference of selection to more frequent genes (or “rich get richer effect” (Elhanati et al., 2014)) on both generated and experimental data (Supplementary Figure 5). Information metrics also did not show any particular selectional trends (Supplementary Figure 6).

Interestingly, while no “rich get richer effect” were observed for the genes frequencies before and after selection, it was detected in terms of generation probabilities (pgens) of CDR3β of DP and SP thymocytes calculated via OLGA model (p < 10^-5^, Mann Whitney test, differences of median log2 *Pgen* between SP and DP was 2.24, Supplementary Figure 1G).

### Comparing TCRα naive repertoire sequencing data to OLGA model generated data

Analysis of α chain CDR3s in the same manner as β chain revealed some common selection effects for both chains along with notable differences. For this analysis a subsample of CDR3 singletons from PBMC (Heikkilä et al., 2021) was taken.

Single amino acids usage analysis revealed a difference in selection processes between α and β chains: no evidence of strong negative selection for Proline and positively charged amino acids was detected. On the contrary, structurally simple amino acids such as Alanine and Serine were negatively selected (Supplementary Figure 7A). Notably, for thymocytes data single amino acid usage trends were barely reproduced (Supplementary Figure 8A).

On a 3-mers level we discover the same trend to remove Cysteines and glycosylation sites during selection. However, the power of this effect was smaller - the median log2 fold change for glycosylation sites in the alpha chain was -0.97, which is 3 smaller than the same value for beta chain (Supplementary Figure 7B). These effects were also observed for thymocytes’ TCRs α chains and it also was weaker than for β chains of thymocytes’ TCRs (Supplementary Figure 8B). None of the putative sulfation sites were affected by selection on both modeled and real world data (Supplementary Figure 7, Supplementary Figure 8)

Unlike the β chain, there was no shared direction of Kidera factors changes for TCRα VJ pairs for both generated and experimental datasets (Supplementary Figure 7C, Supplementary Figure 8C). The absence of common direction of changes could be explained by a larger amount of V and J α genes. We suggest that such diversity may impose the need for VJ-specific selection of the α chain.

The changes observed in the physical properties of TCRα were not identical to those seen in TCRβ. Specifically, the decrease in charge was less pronounced for TCRα than for TCRβ (Supplementary Figure 7D, Supplementary Figure 8D) for both generated and thymocytes data. Similar to modeled CDR3β the winsorizing of modeled CDR3α lengths were also detected (Supplementary Figure 7E). Interestingly, in terms of length, the results obtained for thymocytes TCRα data mismatched with the model-generated one in the same way as it was for TCRβ - shortening instead of winsorizing occurred for thymocytes (Supplementary Figure 8E). Unlike CDR3β, CDR3α’ hydrophobicity remains unchanged after selection in both generated and real world data (Supplementary Figure 7F, Supplementary Figure 8F).

The clusters of most enriched naive T-cells CDR3α sequences shared the same feature with the naive T-cells CDR3β enriched sequences - flexible structure in the middle of CDR3 (such as GGS) (Supplementary Figure 9). This observation is in agreement with the 3-mers analysis where we observed no depletion in GGG 3-mer frequency after selection (Supplementary Figure 8B, Supplementary Figure 9). In thymocytes SP-enriched sequences we observed the abundance of DS and NY motifs (Supplementary Figure 10) instead of simpler ones motifs mentioned above. Depleted clusters demonstrated the abundance of SS 2-mer, which was not detected in relative clusters for the β chain, but presented in enriched TCRα clusters for SP thymocytes (Supplementary Figure 9, Supplementary Figure 10). The role of this 2-mer is rather unclear, however, since they were supported by in-silico and real-world data, we can suggest that they may play an important role in T cells’ biology.

The SoNNia model allowed us to additionally detect increased probability of V amino acid in the third position (Supplementary Figure 4B) in modeled data. However, this effect might be caused by a biased generation model for alpha chains, since we did not observe it for thymocytes data (Supplementary Figure 4D).

Gene usage analysis again demonstrated little or no “rich get richer” effect and barely reproduced the results for real-world data (Supplementary Figure 11, 12). However, the small but significant effect was detected again for generation probabilities calculated via OLGA (Supplementary Figure 8G). Difference between median log2(pgen) of SP and DP was 0.62.

We speculate, that difficulties in catching selection effects on a scale of gene usage for both α and β TCR’s chains may arise from several aspects: (i) different biases of TCR sequencing protocols toward gene usage (Barennes et al., 2021), (ii) flaws in generation model, which mismatches with thymocytes data is especially notable for TCRα, possibly, due to not-sufficient amount of sequences available in open data to train the model (Huang et al., 2022) (Supplementary Figure 11), (iii) different HLA alleles of donors, which samples were taken into analysis (Ishigaki et al., 2022).

Although no clear trend was detected, we suggest that a more in depth-analysis considering such factors as allele similarities, V allele-related TCR expression differences and genomic organization of the TCR locus along with elimination of mentioned above difficulties could be sensitive enough to detect certain selectional effects.

### Single-cell analysis reveals the CDR3 dependent differentiation

Since enriched clusters of CDR3α and β from thymocytes repertoire did not always followed the trends observed in generated data, we adjusted our analysis with single-cell data, which includes both CDR3α and β sequences from PBMC of a healthy donor (https://www.10xgenomics.com/resources/datasets/20-k-human-pbm-cs-5-ht-v-2-0-2-high-6-1-0), to further understand the biological role of these enriched CDR3.

We matched the enriched CDR3 sequences with CDR3 sequences from single-cell data (Fig. 2A, 2B) and annotated cells, which carried matched CDR3s. Cells with TCRs enriched in CD8+ thymocytes sample were expectedly abundant mostly in CD8+ cluster (p < 0.05, Fisher exact test) (Fig. 2A, Fig. 2C). However, CD4+ cells also carried enriched TCRs, highlighting the partial stochasticity of lineage commitment in the thymus and the impact from different MHC alleles. (Yates, 2014).

**Figure 2.**
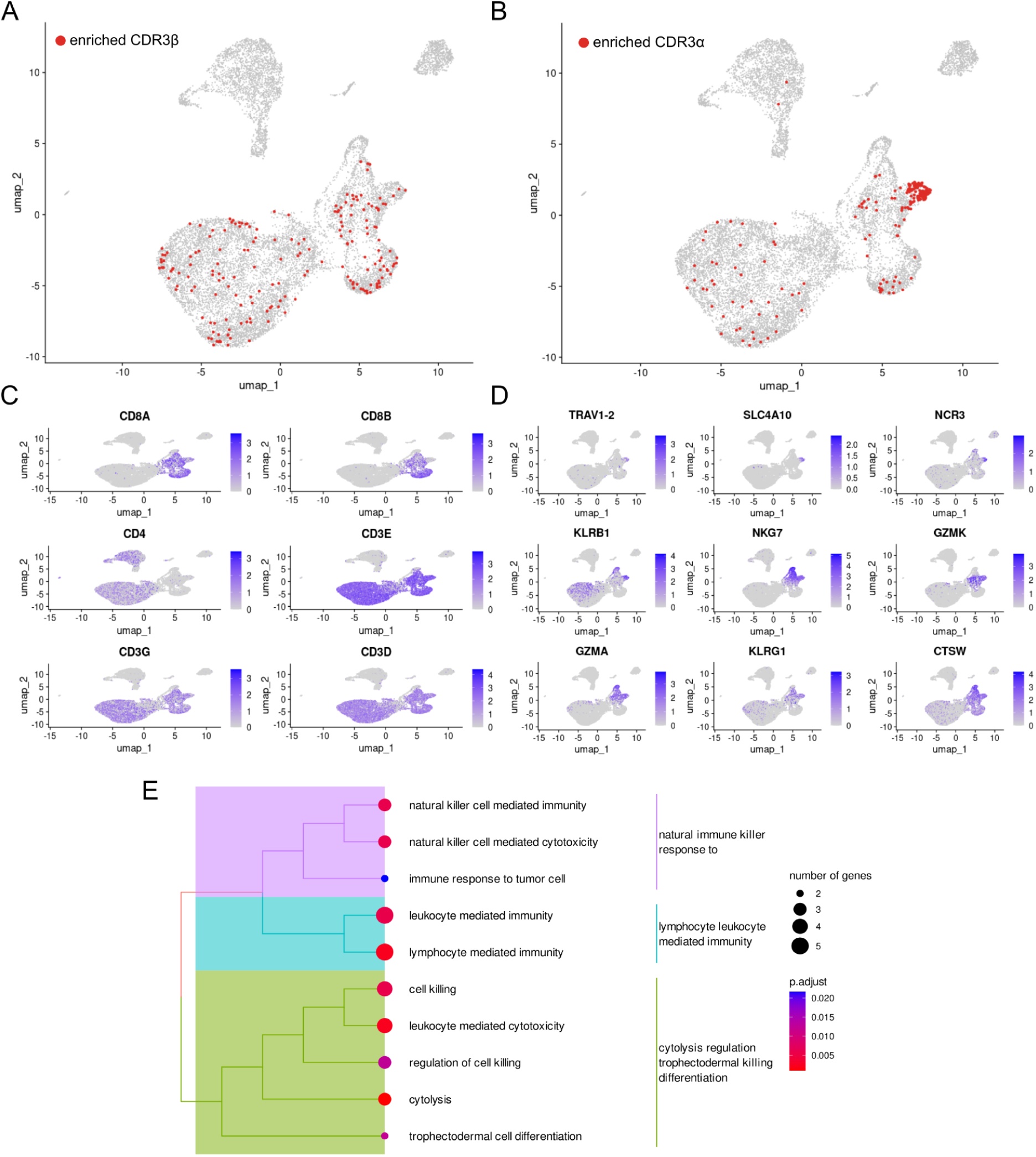
Single-cell analysis of cells carrying enriched CDR3s. **A.** CDR3β enriched repertoires mapped to single-cell UMAP plot. **B.** CDR3α enriched repertoires mapped to single-cell UMAP plot. **C.** T-cell markers on a single-cell UMAP plot. Large proportion of the cells from figure 2A are in the CD8 rich area. **D.** Top 9 differentially expressed genes in cells with CDR3 identical to those enriched in CDR3α CD8 dataset mapped to UMAP plot. TRAV1-2 and KLRB1, which were differentially expressed, are markers of MAIT cells. **E.** Gene ontology (GO) terms enriched in top 20 differentially expressed genes of CDR3α group. They are similar to GO terms enriched in genes expressed by MAIT cells.

For clusters enriched in CDR3α SP sample we discovered a subset of cells densely located in a localized region of the UMAP plot (Fig. 2B). All cells which carried enriched TCRs were taken into differential expression analysis. Visualization of top 9 differentially expressed genes is shown on Fig. 2D. Genes such as TRAV1-2 and KLRB1, which were differentially expressed in our group identify a distinct subset of lymphocytes known as mucosal-associated invariant T cells (MAIT cells) (Godfrey et al., 2019). Such results could be expected since MAIT cells are characterized by their “semi-invariant” TCRs α chains. Additionally, *MR1* which plays role of this cells selection is much less heterogeneous than canonical MHCs (Treiner et al., 2003, Rozemuller et al., 2021). We also repeated this analysis in two other datasets and obtained similar results (Supplementary Note 1).

Furthermore, we examined the cell type TCR abundance in single-cell PBMC data derived from 88 healthy donors. Each enriched or depleted TCR cluster from Supplementary Figures 2, 3, 9 and 10 (every subset of enriched or depleted CDR3 clusters which was analyzed above) was intersected with single-cell TCR repertoire. Then the abundance of these clusters in a particular cell type was tested. Note that we used only the top 5 largest clusters in each enriched or depleted subset as it described in methods.

Generally, we received an expected picture of TCRs distribution among cell types - TCR from depleted clusters were barely found in any cell type subset while TCR from enriched clusters were found mostly in naive cells (Fig 3). SP thymocytes clusters were mostly represented by CD8+ naive cells, which is expected since SP thymocytes in our research are of CD8+ lineage (Fig. 2, 3B). The numbers of TCRs from depleted clusters found in Single Cell data were 10-20 smaller than from enriched clusters (Fig 3 C, D, G, H). Latter indicates that depleted clusters are a good proxy of TCRs which will not pass the thymic selection.

**Figure 3.**
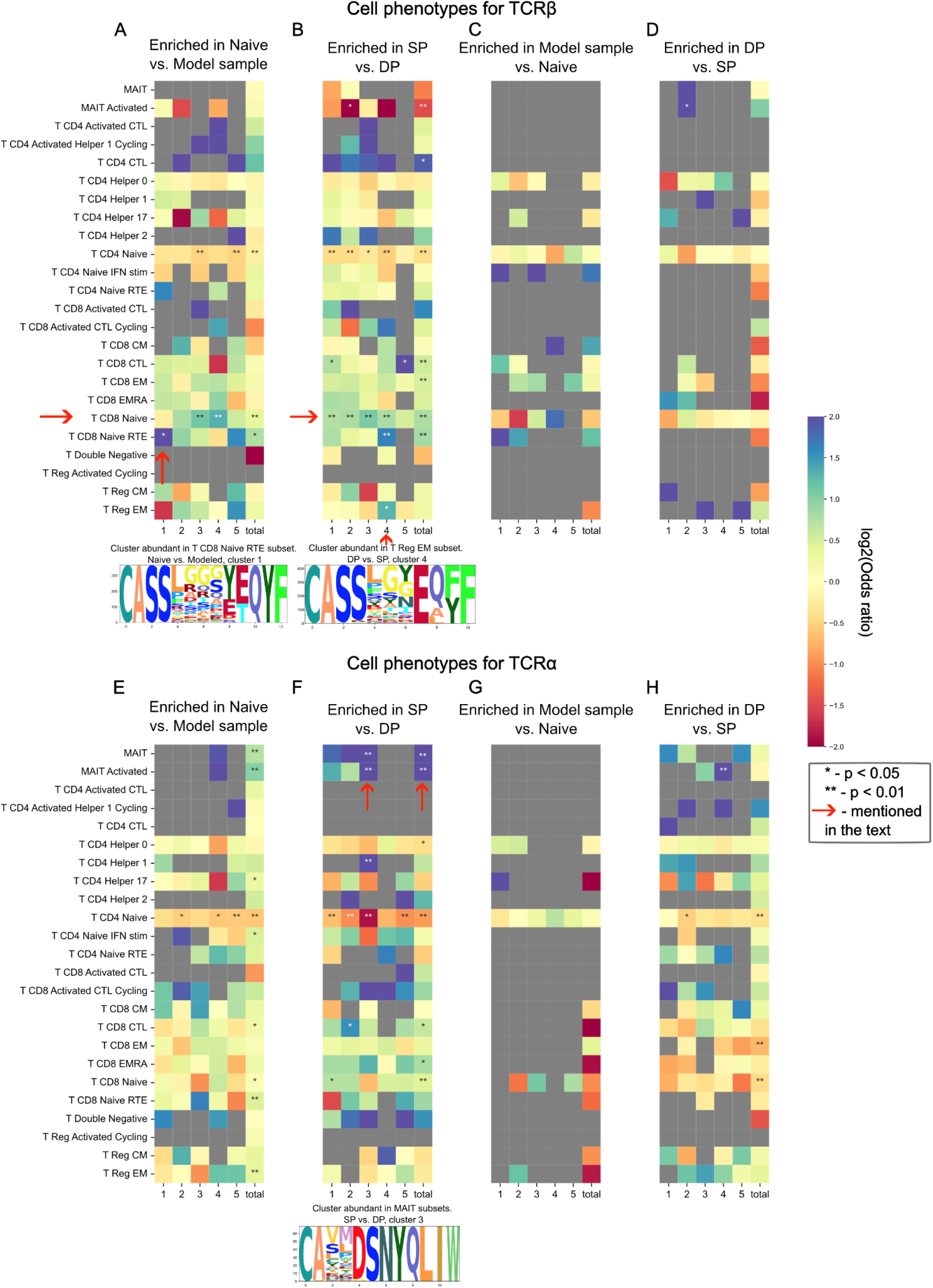
Abundance of TCR sequence motifs enriched and depleted during thymic selection in T-cell subsets defined by single-cell sequencing data. Odds ratio of observed to expected number of sequences matching between cell phenotype and selection “motif” (cluster) is shown by color, two-tailed Fisher exact test P-values for odds scores post multiple testing correction are shown with asterisks (* for p < 0.05 and ** for p < 0.01). Cluster-phenotype pairs discussed in the main text are highlighted with arrows. Chosen clusters are represented as logos. SP, DP, Naive and Model datasets for TCR alpha and beta are shown in panels **A-H** represent different comparisons as described in the main text: **ABEF** represent negatively selected TCRs, while **CDGH** represent positively selected ones; **ACEG** are based on model data and naive cells, **BDFH** are based on real data; **ABCD** describe beta chain, **EFGH** describe alpha chain.

Another interesting observation is considering enriched after selection naive TCRβ clusters. It appears that TCRβs of CD8+ cells are more prone to neighborhood enrichment than TCRβs of CD4+ cells (Fig. 3A). Additionally, cluster enriched in CD8+ Recent Thymic Emigrants (RTE) subtype stood out against the other clusters enriched in CD8+ Naive cells (Fig. 3A). Such observation could be the sign of peripheral selection effects on the repertoire. Cluster 4 of SP thymocytes is also of particular interest since it demonstrated abundance in both CD8+ TRE and Tregs (Fig. 3B).

Effects linked with MAIT cells were also traced in this analysis. We discover strong enrichment in the MAIT cells subtype for the cluster 3 of TCRα enriched in SP thymocytes (Fig. 3F). Such an effect was already detected in previous analysis (Fig. 2).

### Structural analysis supports observed differences in CDR3s of CD4 and CD8 repertoires

Next, we compared CD4+ and CD8+ naive cells’ TCR repertoires (Qi et al., 2014). The most notable result of this comparison is clusters enriched in CD4+ or CD8+ CDR3s. We employed the TCRmodel tool to analyze the structure of the most prevalent CDR3 in the largest clusters within each group (Yin et al., 2023) and discovered a difference in convexity between enriched CDR3 CD8+ and CDR3 CD4+ structures. (Fig 4A, B). It appeared that CD4+ enriched clusters were structurally flat, whereas CD8+ clusters were more convex.

**Figure 4.**
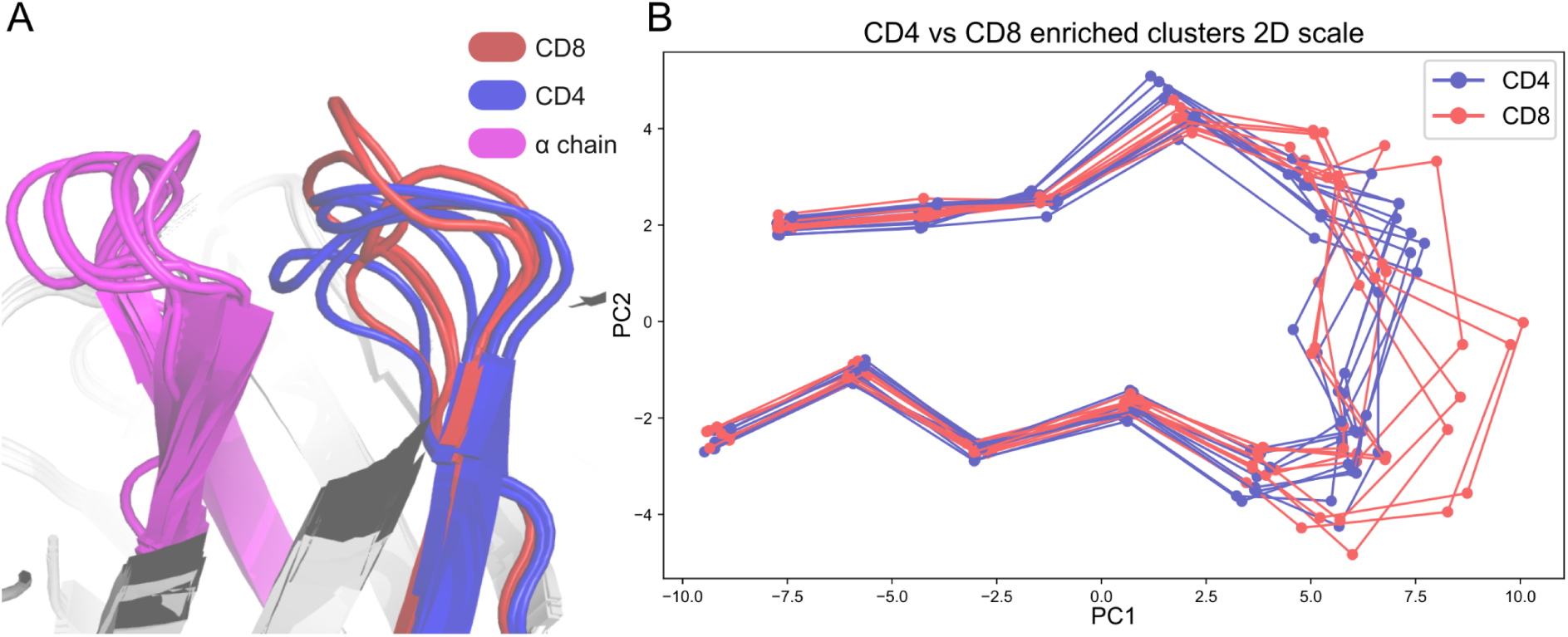
Structural differences between CDR3 of CD8+ and CD4+ T-cells. **A.** Modeled 3D structures of CD4 and CD8 enriched CDR3β repertoires. CD8 enriched structures tend to be more loose while CD4 enriched clusters are more assembled. **B.** Same as A but 2D PCA projection with additional sequences from the same clusters with 2 additional TCR for each cluster.

This fact may be explained by the conformation of peptides in MHC class I and MHC class II grooves. MHC class I, which recognised by CD8+ cells, tends to present peptides with a middle bulge, whereas MHC class II, recognised by CD4+ cells, tends to present flat peptides (Cole, 2013).

### HLA allele affects the selection

Next, we considered the HLA alleles influence on the thymic selection. It is still a subject of debate whether HLA alleles affect the selection or not (Isacchini et al., 2024).

We used CDR3s bulk sequencing data of 3 pairs of twins aged 20 to 23 to answer this question (Pogorelyy et al., 2018). This data contains TCRβ repertoires for 3 pairs of twins (assigned as S, P and Q) in 2 replicas - day of vaccination before the shot (day 0) and a day prior to vaccination day (pre-day).

We compared their naive CDR3 clusters to modeled background repertoire. On the one hand, we expected their enriched naive CDR3 clusters to be closely related with each other since twins from the same pair have the same HLA alleles. On the other hand, we anticipated twins from different pairs to be different from each other in context of their naive TCR repertoire.

We extracted the same number of CDR3s from each twin sample and then clustered the enriched sequences from each sample together. Having this done, we obtained a number of clusters composed of sequences from different samples. Supposedly, each cluster contains CDR3s which are functionally similar to each other (recognise similar peptides in the MHC-peptide complex). Thus, we would anticipate CDR3s derived from the same pair of twins to fall into the same clusters.

To verify this hypothesis, we analyzed the proportion of CDR3s from a particular twin in the clusters obtained. In this representation, each sample can be viewed as a discrete probability distribution of appearing in a particular cluster. This allowed us to calculate Jensen–Shannon divergence between samples (Fig. 5A). We discovered that samples within each twin pair are closer than samples from different twin pairs. Such proximity can also be observed in tSNE plot (Fig. 5B).

**Figure 5.**
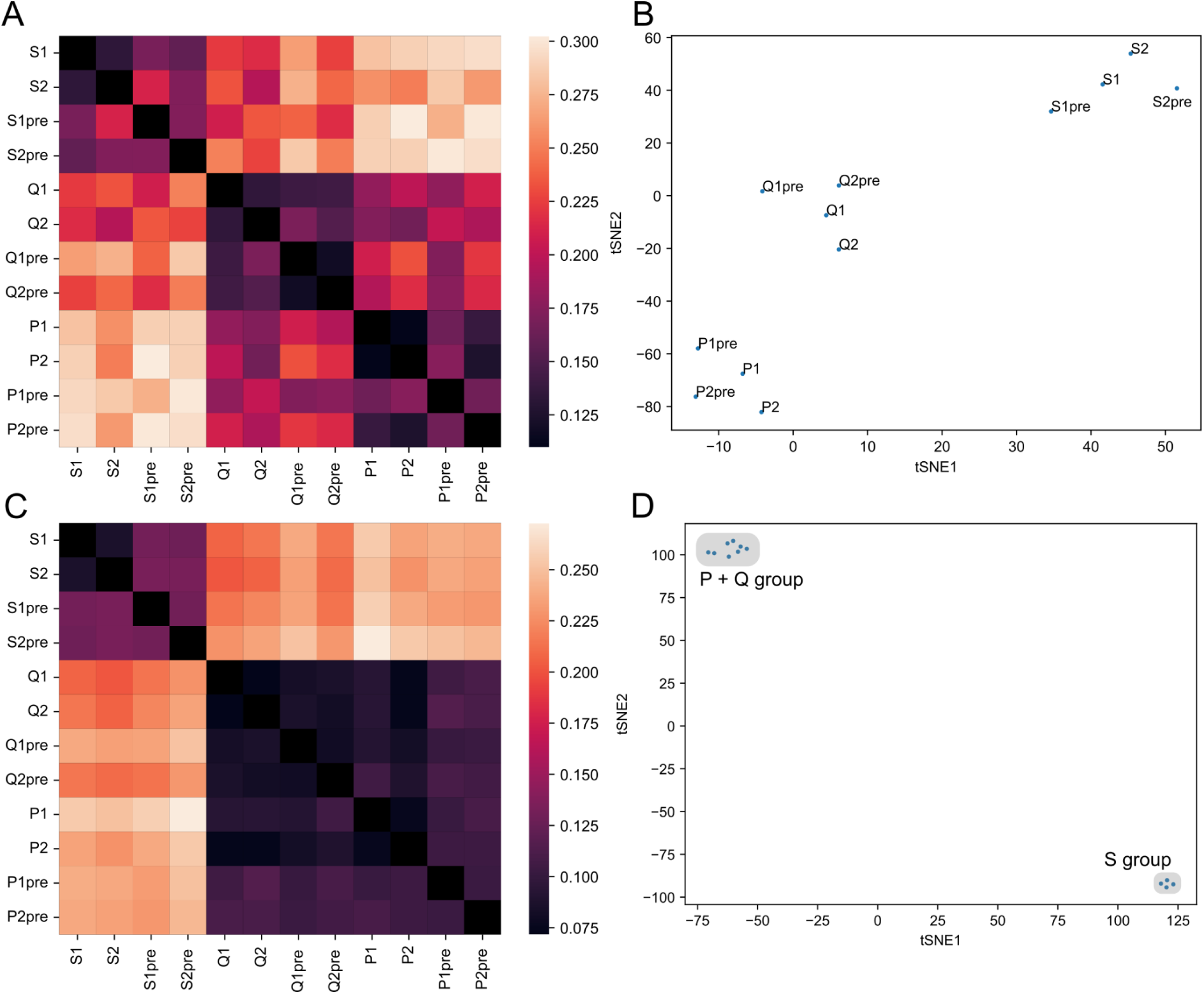
HLA allele affects thymic selection of the TCRβ chain. **A.** Heatmap representing Jensen-Shannon divergence between frequencies of occurrence in enriched functional clusters for each twin sample (*S*, *P* and *Q* pairs in 2 replicas). **B.** t-SNE plot for occurrence frequencies in positively selected functional clusters. **C.** Heatmap representing Jensen-Shannon divergence between frequencies of occurrence in modeled data functional clusters with each twin sample as a background. **D.** t-SNE plot for occurrence frequencies in negatively selected functional clusters.

Notably, twin pairs with the homozygous HLA-A 02:01 allele (pairs P and Q) appeared closer to each other than to the S pair which had only one copy of HLA-A 02:01 allele. HLA haplotypes of taken donors can be found in Supplementary Table 1.

The analysis of the clusters which were depleted during the selection showed similar results. (Fig. 5C, Fig 5D). Thus, HLA alleles do not only affect TCRs promoted by selection, but also the TCRs which are eliminated by it, so called “holes” in repertoire. We detected the effect of homo/hetero zygosity of HLA-A gene allele on the T-cell repertoire again but this time it was more pronounced. In terms of depleted clusters, twin pairs, which were homozygous by HLA-A (P and Q), were 2-3 times closer to each other than to heterozygous by HLA-A twin pair (S). (Fig. 5D).

We repeated the analysis in the same manner for 2 other sets of twins’ TCRs. One of these sets consisted of CD4+ naive cells. We obtained similar results or showed that the sample size was not sufficient to carry out the analysis (Supplementary Note 2).

These results suggest that one can observe the imprint of HLA-based selection on the naive repertoire. Moreover, this imprint was observed in both enriched and depleted clusters, indicating that particular HLA alleles are favorable for one TCRs and unfavorable for another. Thus, genetic diversity of HLA alleles results in diverse TCR repertoires in the population.

## Discussion

In this study we sought to identify and investigate factors which are crucial for passing thymic selection. We considered such factors as amino acids composition, K-mers composition, Kidera factors, charge, length, and hydrophobicity, TCR gene usage, along with analysis of functional clusters enriched in samples before and after selection and SoNNia inferred marginal probabilities of amino acids to occur in a particular position of CDR3. To gain a deeper understanding of enriched clusters additional means such as analysis of physical structures and single-cell data analysis were harnessed. Finally, we considered HLA alleles as a factor which has an impact on selection and indeed managed to show a significant influence from it.

Amino acids usage analysis revealed that having Proline and large positively charged amino acids in CDR3 reduces the likelihood of TCRβ survival during selection (Fig. 1A, Supplementary Figure 1A). However, we found no such effects on the survival of CDR3α (Supplementary Figure 7A). On a 3-mers level we showed that Cysteine and N-glycosylation sites negatively affected the chances of survival (Fig. 1B, Supplementary Figure 1B, 7B, 8B). Kidera factors analysis revealed a difference in selection effects for α and β CDR3s. For CDR3β, we observed synchronous changes in 4 Kidera factors for every VJ pair (Fig. 1C).

These 4 Kidera factors represent physical volume, hydrophobicity, and occurrence in the alpha region. Thus, small, hydrophobic, and Glycine-reach CDR3βs have greater chances of surviving selection. Unlike CDR3β, we did not observe the common direction of changes for the CDR3α Kidera Factors for every VJ pair (Fig. Supplementary Figures 7C, 8C). This may indicate that physical properties selection in the case of CDR3α is specific to the VJ pair. At a physicochemical level, we observed a decrease in the charge and length of the CDR3α and CDR3β repertoires after selection (Fig. 1D, Supplementary Figures 1E, 7D, 8E). However, there were differences in hydrophobicity changes after selection between α and β TCRs. More hydrophobic CDR3s β were likely to survive (Fig. 1F). In contrast, CDR3α’s hydrophobicity did not influence survival (Supplementary Figure 7F, 8F). Such inconsistency may arise from VJ specific physical factors selection in the case of CDR3α. Cluster enrichment analysis revealed that flexible CDR3 with poly-Glycine subsequences tend to survive the selection and this effect is more pronounced for β CDR3s. Furthermore, structurally complex CDR3s tend not to pass the selection (Supplementary Figures 2, 3, 9, 10). Gene usage analysis demonstrated a vague picture of gene usage changes through the selection (Supplementary Figures 5, 6, 11, 12).

By supplementing our analysis with single-cell datasets with adjusted CDR3 sequencing data, we showed the direct influence of CDR3 on T-cell lineage commitment (Fig. 2, 3). This was especially notable for CDR3α. CDR3α is mostly responsible for commitment to specific non-conventional lineages such as MAIT cells, which CDR3s were strongly enriched in our analysis (Fig. 2)(Godfrey et al., 2019). Structural analysis of enriched CDR3β revealed that commitment to CD4/CD8 lineages is mostly influenced by CDR3 physical structure (Fig. 4). Cells carrying structurally flat CDR3s tend to commit to CD4 lineage since peptides represented by MHC class II have flat structure. In contrast, CD8+ T-cells’ CDR3s are structurally curved similar to peptides presented by MHC class I (Cole, 2013). Finally, we demonstrated the effect of HLA alleles on thymic selection using CDR3 repertoires from 3 pairs of twins. This result contradicts previous reports (Isacchini et al., 2024).

Overall, our analysis showed that structurally simple and flexible CDR3s have a greater chance of passing the selection. We hypothesize that such CDR3s are capable of recognizing a wide variety of peptides within the thymus with a moderate binding strength. This ability meets the exact requirement for getting through positive and negative selections (Ashby and Hogquist, 2023). However, it is important to note that there is no consensus considering polyreactivity of maturing T-cells within the thymus. (Yates, 2014).

While most of our findings can be explained by the aforementioned paradigm some of them remain rather strange, in particular the difference in selection preferences for CDR3α and CDR3β. The presence of non-conventional T-cells, such as MAIT cells, in our analysis may partially explain this contradiction. These cells are selected in the thymus by the different mechanisms, mostly driven by the α chain (Godfrey et al., 2019).

The difference in amino acid preference is peculiar since no effect on survival from presence of Proline and positively charged amino acids in CDR3α were observed. However, these amino acids will have a strong impact on TCR structure. It is also unclear why the hydrophobicity which had a strong impact on CDR3β selection did not affect CDR3α as well. However, we demonstrated that hydrophobicity of CDR3α repertoire was already comparatively high even before selection. The effect of glycosylation site incorporation in CDR3 is also weaker for CDR3α. Finally, on the level of VDJ recombination generation probabilities the effect of selection for β chain was 3.6 times larger. The possible explanation for these inconsistencies may be in the presence of rescue mechanisms for an α chain. While CDR3β has only two attempts to form a functional receptor and it undergoes additional β selection the CDR3α can undergo rearrangement several times during functional TCR formation (Duan and Mukherjee, 2016). Also, it is important to note that, as we observed in this study, the choice of V and J genes plays a crucial role for an α chain selection process but not for β chain. Finally, the observed negative impact of SS dimers on survival of TCRα is still enigmatic.

In this paper, we demonstrated that probabilistic model-generated data can be utilized as a model for unselected TCRs. However, there are some limitations. First, the difference in protocols for obtaining data for the model training and for the data used in our study might explain the difference in results obtained from the in-silico and real-world data (Barennes et al., 2021; Bolotin et al., 2012; Sethna et al., 2019). Particularly, the absence of “winsorizing” of CDRs’ lengths in real-world data and different trends for gene usage between thymocytes and generated data could be explained by such inconsistency. Second, It is still an open question how many TCRs should be generated for robust analysis. Notably, selection trends for generated TCRα were more often in disagreement with real world data than ones observed in generated TCRβ repertoire. We might speculate that this is because of the lack of TCRα data generally available to train the OLGA/IGoR model used in this study (Huang et al., 2022). Nevertheless, the results obtained using in-silico data are mostly in agreement with results obtained in real-world data.

It is important to note that probabilistic model generate TCR repertoire which was not a subject of any functional selection. However, double positive TCRs, which were used as a relative sample to model-generated repertoire, had already undergone β selection, during which some unfunctional CDR3s were eliminated. Thus, these two samples are not entirely biologically comparable. (Dupic et al., 2019, Li et al., 2021).

Singletons’ approach for simulating naive T-cells has its own shortcomings too. While most of CDR3s indeed represented naive lineage some of these cells may still be of different lineages (Britanova et al., 2014).

Single positive thymocytes, which were used in our study alongside naive cells, did not fully represent this cell type, since they still can be negatively selected in the thymus medulla in the late stages of thymic selection (Duan and Mukherjee, 2016). As a result, we may miss certain negative selection effects when using this data.

The approach for CDR3s characterization by matching them with publicly available single-cell data, in our opinion, has a great potential since it allows, in terms of a lack of experimental data, to gain a deeper understanding of the biology of a particular group of CDR3s. However, differences in HLA alleles across donors may impose some restrictions on this analysis. Such restrictions may be the exact reason why we observed dense cluster of enriched TCRαs in the MAIT cells’ part of the UMAP plot, although enriched TCRs β demonstrated a diffuse distribution among various cell types with only a bias toward CD8+ cells. (Fig. 2). The possible reason of this inconsistency may lay in the fact that while MAIT cells are selected on a *MR1* gene which is highly conservative among individuals (Treiner et al., 2003), the MHC I and MHC II show much greater allele diversity (Markov and Pybus, 2015). However, this difference may be at least partially explained by the stochastic nature of lineage commitment (Yates, 2014). The software for TCR structure modeling is also very helpful in terms of very little structural data available (Yin et al., 2023). Overall, we demonstrated that CDR3s significantly affect the T-cell lineage commitment using structural and single-cell analysis.

Comparative analysis of functional CDR3 clusters allowed us to reveal an HLA allele-mediated effect on TCRs selection. This result contradicts prior findings (Isacchini et al., 2024), which suggested an overall lack of forbidden sequences for the selection, thus neglecting among others the HLA-alleles impact on the process of thymic selection. However, the suggestions of possible impact of HLA allele on selection were made before (Ishigaki et al., 2022). Moreover, similar study design, involving 6 twin pairs with paired TCRαβ sequences available, were used to demonstrate influence of genetic factors on TCR repertoire (H et al., 2020). Disagreement with previous results may have roots in the difference of approaches being used. While Isacchini et al. analyzed generation probabilities of nucleotide sequences which can only intermediately be linked with physical nature of peptides recognition, we analyzed the functional landscape of CDR3s. Furthermore, soNNia model in Sacchini et al. study was trained in mixed sample (samples from different subjects were analyzed together) which made it impossible to detect any subject-specific effects in that section of their study. One can note that our results may be biased because of the probable similarities of antigens which were met during co-living of twins from a particular pair. However, this does not explain the observed effect from HLA-A homo/heterozygosity. The HLA repertoire from twins with homozygous HLA-A 02:01 allele were closer to each other than to ones with one copy of HLA-A 02:01 (Fig. 5). Interestingly, it was shown before that CD4+ repertoire from mice homozygous by HLA is more diverse than repertoire from heterozygous ones (Brown et al., 2024). Latter facts one more time highlighted the existence of “holes” in repertoire induced by negative selection. Thus, we can conclude that there are indeed forbidden and favorable for selection sequences, which are not determined by generation probability alone.

Consideration of HLA alleles mediated effects goes beyond thymic selection. For example, it was shown that patients with particular HLA alleles tend to respond differently to treatment with immune checkpoint inhibitors (Chowell et al., 2018). The possible effect of HLA alleles in cancer treatment was also suggested by Schumacher et al. (Schumacher et al., 2019). Moreover, it was proposed that HLA may play a role in side effects of vaccines (Bolze et al., 2022). Finally, particular HLA alleles combinations are one of the most noticeable risk factors of type I diabetes (Noble and Valdes, 2011). Altogether, the study of HLA alleles-mediated effects can lead the way to better personalized medicine and deeper understanding of particular diseases’ pathogenesis.

Overall, our results may have a number of practical applications. Using the set of properties which were obtained in this study one can assess the likelihood of passing the selection for the particular TCR. This may be used, for example, in a study of autoimmunity, since autoimmune clonotypes should have a very low chance of passing the selection. More broadly, our findings can form the basis for machine learning models aiming to predict the probability of TCR to reach the naive state or “in-silico thymic selection” model.

Another possible implementation of our results is in the field of immunotherapy. Our findings could be important in the design of adoptive T-cell transfer. This therapy involves the engineering of TCR which recognises desirable antigens (Kalos and June, 2013). Using findings presented in this study one can arrange probable TCR variants to select only those that can mimic natural ones. Paying attention to our results may also improve affinity to antigens making it more physiological. This could decrease the side effects of such therapies. Our results also could be applied in the design of chimeric antigen receptors (CAR). For example, according to our findings, the incorporation of glycosylation sites in CAR should be avoided. This is also true for immune checkpoint inhibitors design and any other therapeutic antibody.

Before our study, glycosylation sites were reviewed only in terms of antibodies. Authors highlight de-novo acquisition of glycosylation as a distinctive property of follicular lymphoma (Zhu et al., 2002). Other authors suggested that levels of N-glycosylation acquisition differ between follicular lymphoma subtypes and generally may have an important role for this malignancy in activation and interactions with other cells (Schneider et al., 2012, Leich et al., 2021). Therefore, we might propose that acquired N-glycosylation may have a diagnostic value in case of T-cell lymphomas as well.

Checking the theories that infrequently recombined TCRs rarely pass the selection process and vice versa, or the “rich get richer” effect, is another essential application of our findings (Elhanati et al., 2014). One can assess the generation probability of the clonotypes with particular features highlighted in this study and compare it with generation probabilities of clonotypes with opposite features.

All things considered, our research identified a number of TCR characteristics that significantly influence thymic selection. These results offer a quantitative explanation of the entire selection process and a clearer picture of the range of potential repertoire feature changes that may take place during thymic selection. It also provides an avenue for additional study and experimental validation, along with potential real-world applications.

Additional thymocyte datasets could be useful to expand and validate the results of this investigation. We also experienced the lack of datasets with paired α and β chains. Although this data is more expensive to generate than bulk rep-seq (Rosati et al., 2017) it could give an opportunity to study the selectional behavior of the whole TCRs heterodimer, allowing to detect mutual α-β effects of selection.

## Acknowledgments

We would like to thank Dr. Dratva and Dr. Kretschmer for their assistance with single-cell data annotation knowledgebase (T-cell phenotypes and receptor sequences from Lindeboom *et al*.). This work was supported by the Russian Science Foundation (RSF) grant №25-75-30013.

## Supplementary Figures and Tables

**Supplementary Figure 1.**
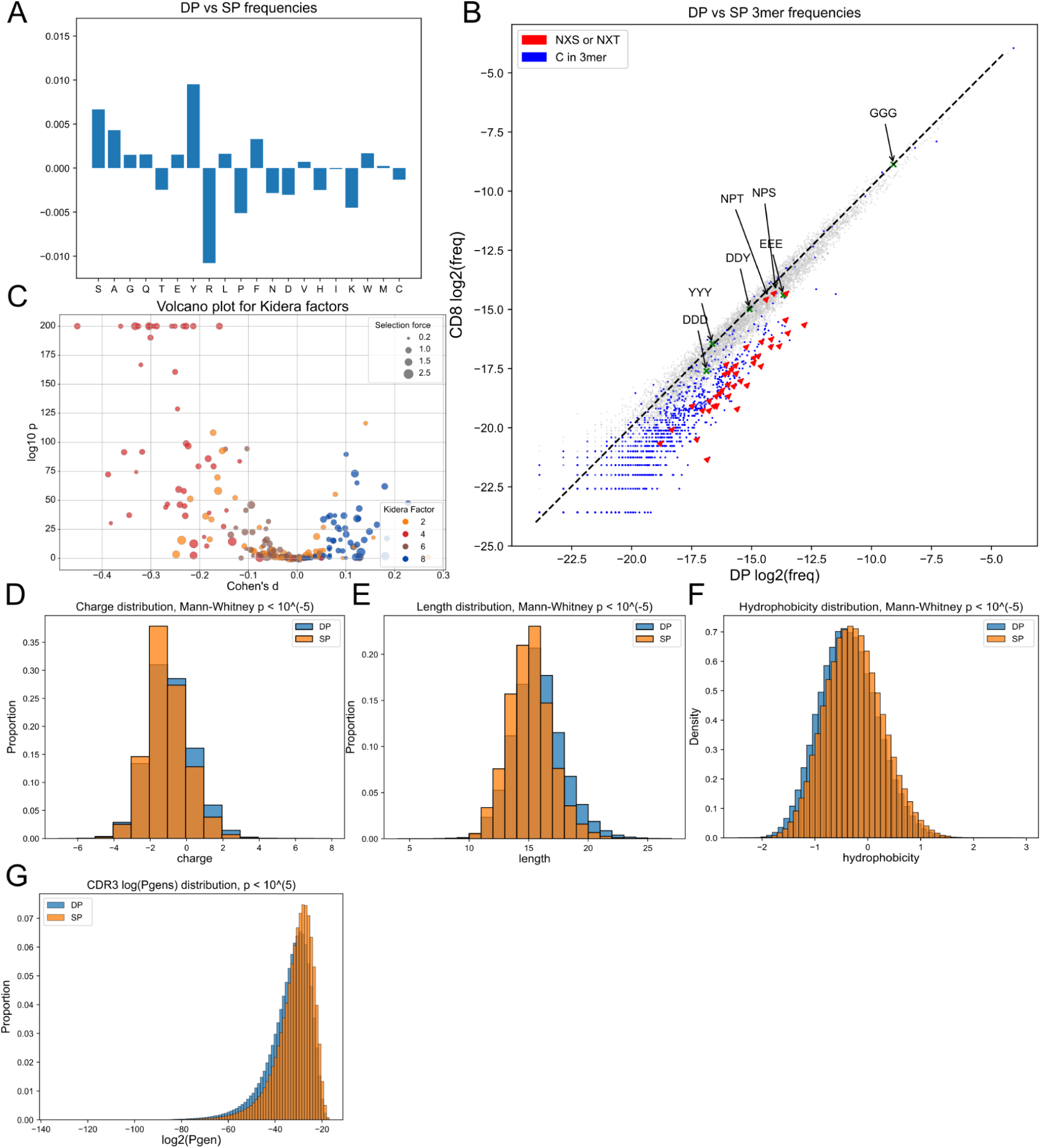
Comparing CDR3β sequences of double positive (DP) thymocytes and CD8+ single positive (SP) thymocytes. **A.** Amino acid frequency comparison between double positive and single positive thymocytes. **B.** Comparing frequencies of k=3-mers between thymocyte subsets. Glycosylation sites (red) and Cys-containing 3-mers (blue) are negatively selected. **C.** Volcano plot showing positive and negative selection P-values and effect size for four selected Kidera factors. Each point represents an independent calculation performed for a certain VJ pair. Results are similar to those observed in data generated using the VDJ rearrangement model, except for the effect for Kidera factor 6 that is smaller. **D.** Negative shift of the charge distribution post-selection. **E.** Shortening of CDR3 lengths post-selection. **F.** Positive shift of the hydrophobicity distribution post-selection. **G.** Generation probability (*Pgen*) distribution is higher for SP thymocytes compared to DP, in line with previous observation that variants with higher *Pgen* are more likely to pass selection.

**Supplementary Figure 2.**
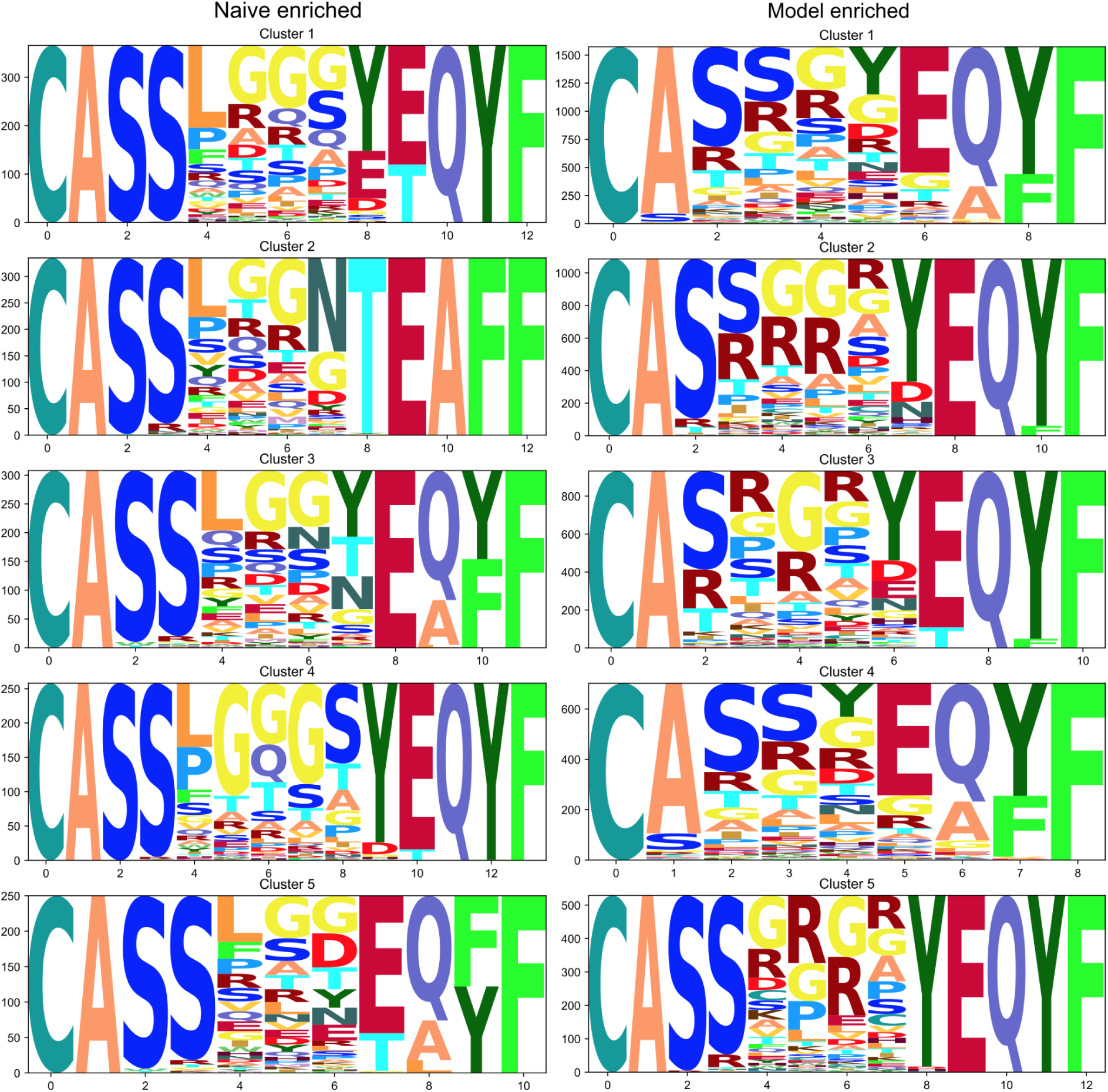
Top 5 largest clusters of TCR-CDR3β sequences enriched in naive T-cell sample (positively selected, left panel) and in a sample generated using VDJ rearrangement model- (negatively selected, right panel). Note that flexible poly-G motifs are favored by selection while Arg residues are negatively selected.

**Supplementary Figure 3.**
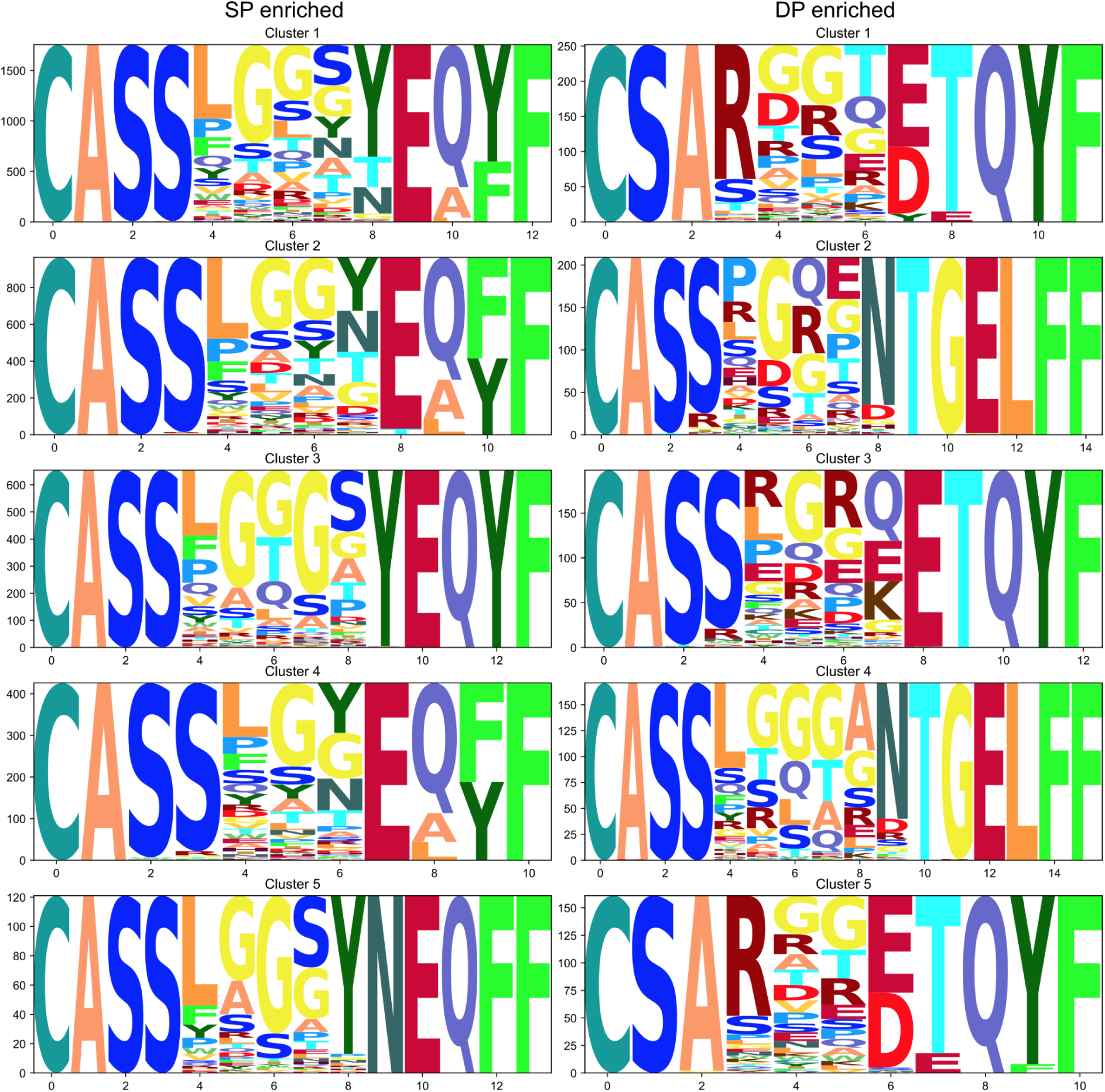
Same as in **Supplementary Figure 2**, but comparing CD8+ single-positive (SP) as a proxy for positively selected TCR-CDR3β motifs and double-positive (DP) as a proxy for pre-selection thymocytes.

**Supplementary Figure 4.**
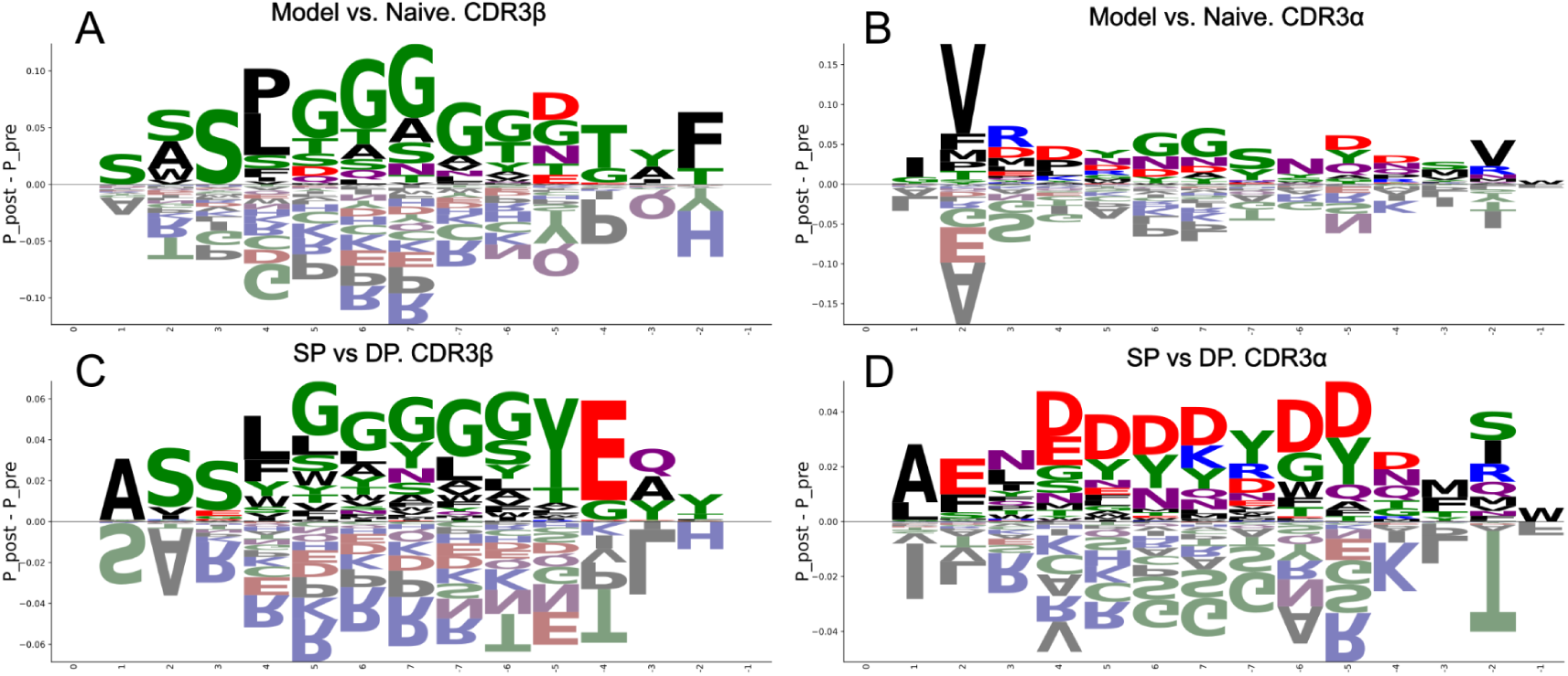
Difference in marginal probabilities of amino acid occurrence learned by SoNNia thymic selection model visualized using sequence logos. VDJ rearrangement model vs naive cells and DP vs SP CDR3α and β are shown.

**Supplementary Figure 5.**
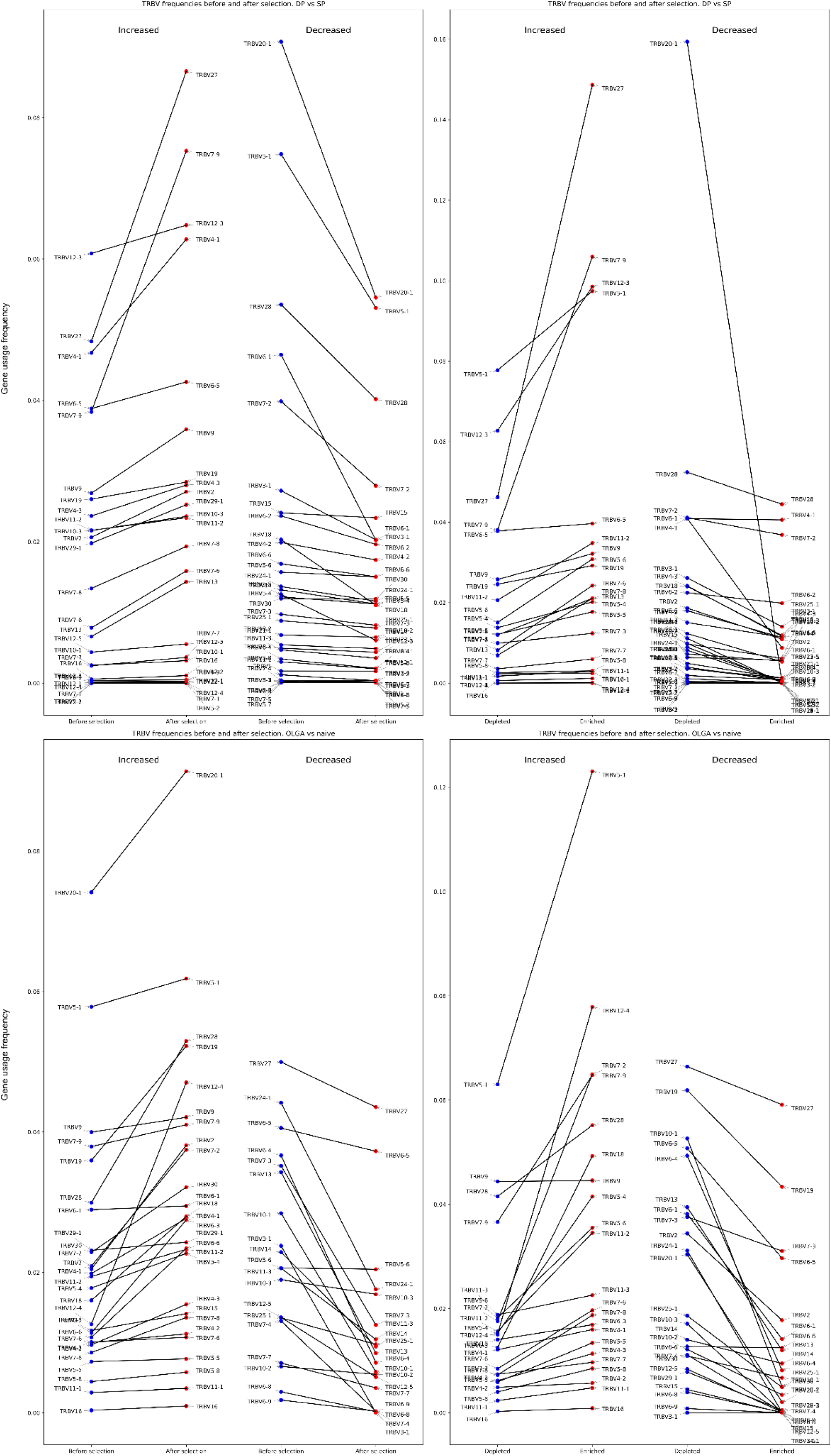
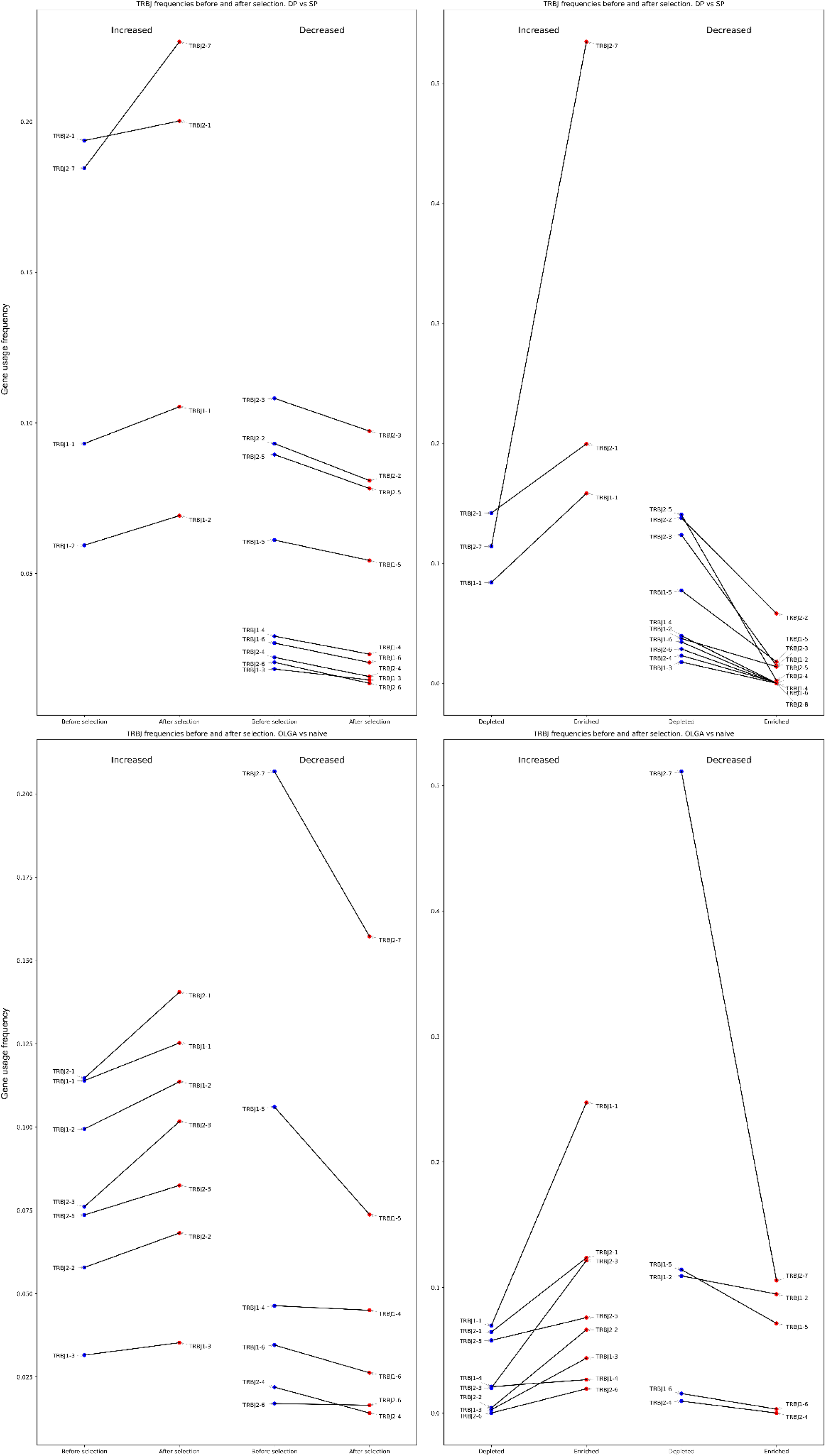
TCRβ gene Variable and Joining gene usage before and after selection. Analysis was carried out both for whole repertoires and for selected enriched and depleted TCRβ CDR3 clusters.

**Supplementary Figure 6.**
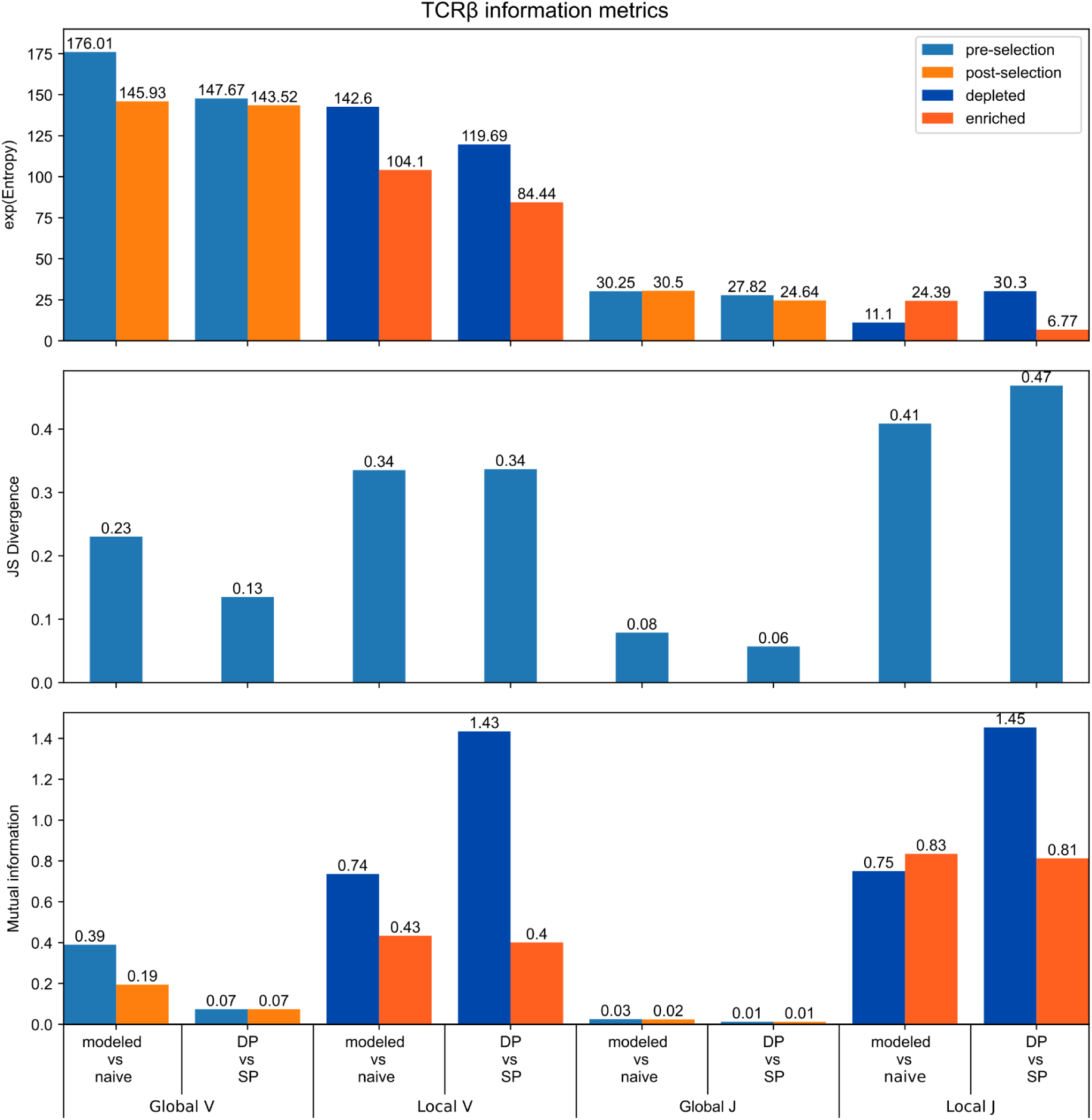
TCRβ V and J gene usage information metrics. Analysis was carried out for a whole repertoire (Global) and for enriched pre-/post-selection TCRβ CDR3 clusters (Local).

**Supplementary Figure 7.**
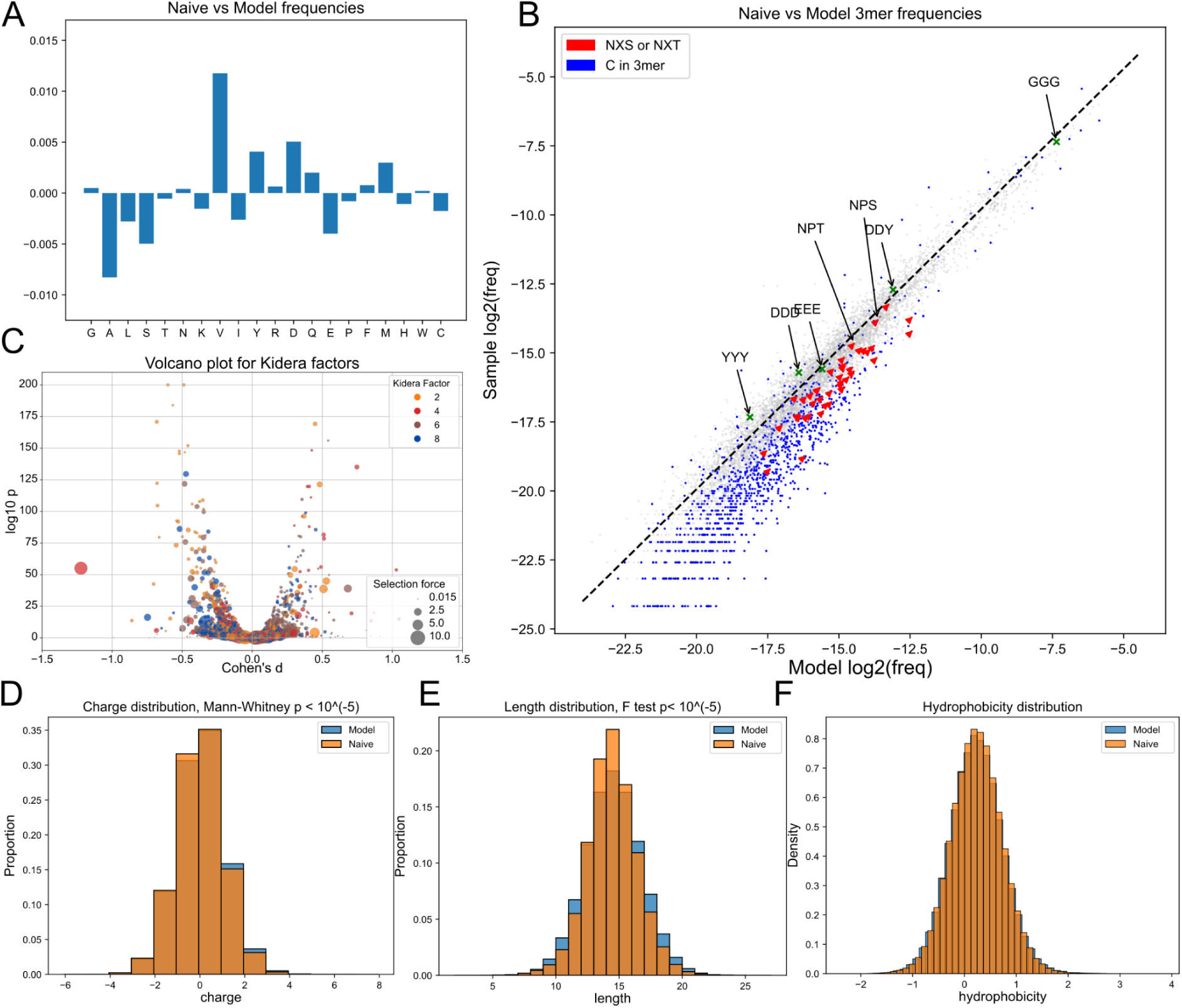
Same as Figure 1, but for the TCRα chain. Model-generated CDR3α and naive CDR3α repertoire comparison. **A.** Amino acid frequency comparison between repertoire generated with VDJ rearrangement model and naive T-cells. **B.** Comparing frequencies of k=3-mers between model generated and naive subsets. Glycosylation sites (red) and Cys-containing 3-mers (blue) are negatively selected. **C.** Volcano plot showing positive and negative selection P-values and effect size for four selected Kidera factors. Each point represents an independent calculation performed for a certain VJ pair. Direction of selection appears to be dependent on VJ-pair choice since there is no common direction for each Kidera factor selection. **D.** Negative shift on charge distribution post-selection. **E.** “Winsorizing” of CDR3 length distribution post-selection, extremely short and long CDR3s are removed during selection. **F.** Hydrophobicity remains unchanged post-selection unlike CDR3β data.

**Supplementary Figure 8.**
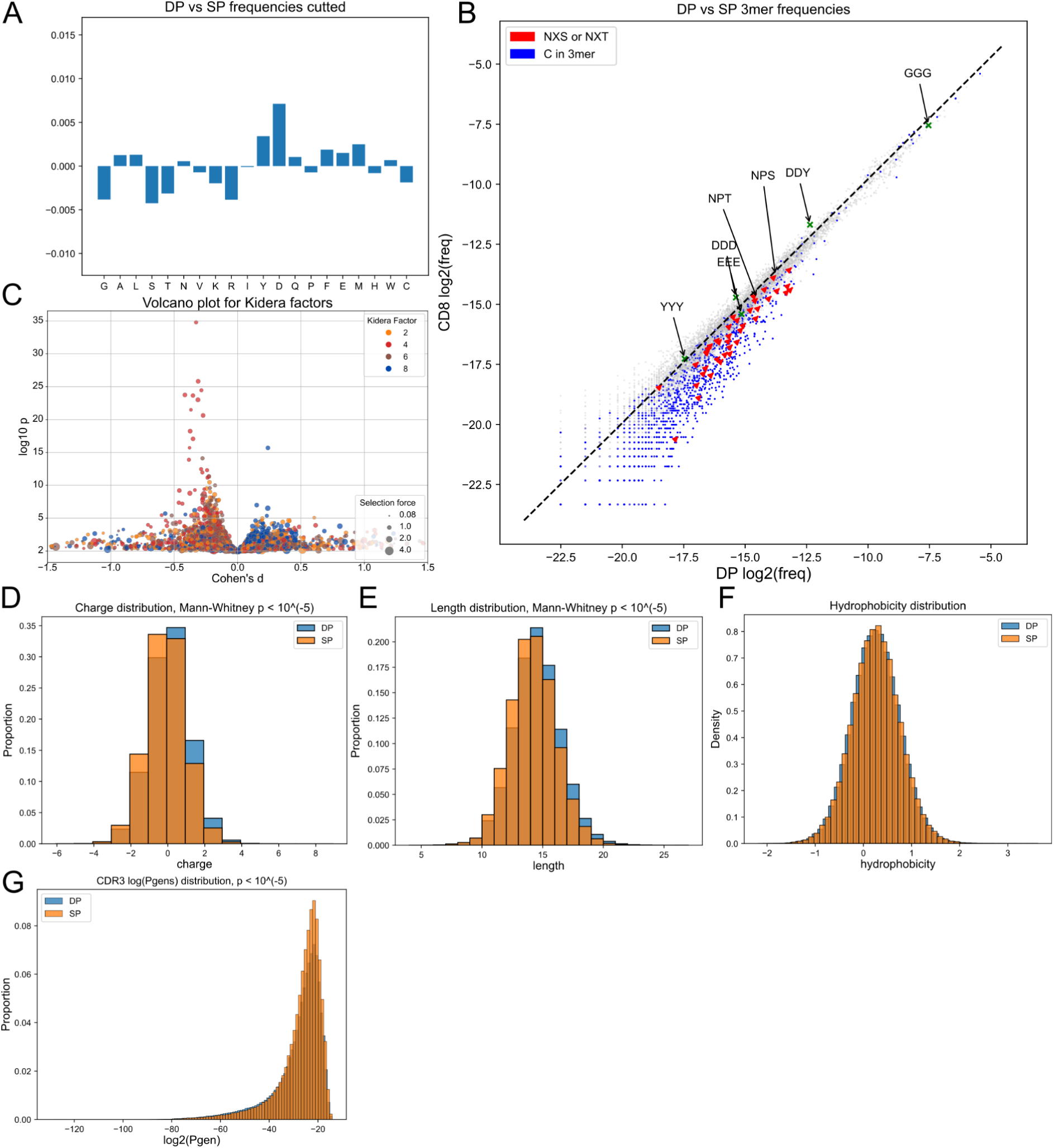
Same as **Supplementary Figure 1**, but for the TCRα chain. Double positive (DP) thymocytes CDR3α and CD8 single positive (SP) thymocytes CDR3α repertoires comparison. **A.** Amino acid frequency comparison between double positive and single positive thymocytes. **B.** Comparing frequencies of k=3-mers between thymocyte subsets. Glycosylation sites (red) and Cys-containing 3-mers (blue) are negatively selected. **C.** Volcano plot for Kidera factors affected by selection. Each point represents a VJ pair. Change appears to be dependent on VJ choice as it was observed on generated data. **D.** Negative shift of the charge distribution post-selection. **E.** Shortening of CDR3 lengths. **F.** Hydrophobicity remains unchanged post-selection as it is in the generated data. **G.** Generation probability (*Pgen*) distribution is higher for SP thymocytes compared to DP, in line with previous observation that variants with higher *Pgen* are more likely to pass selection.

**Supplementary Figure 9.**
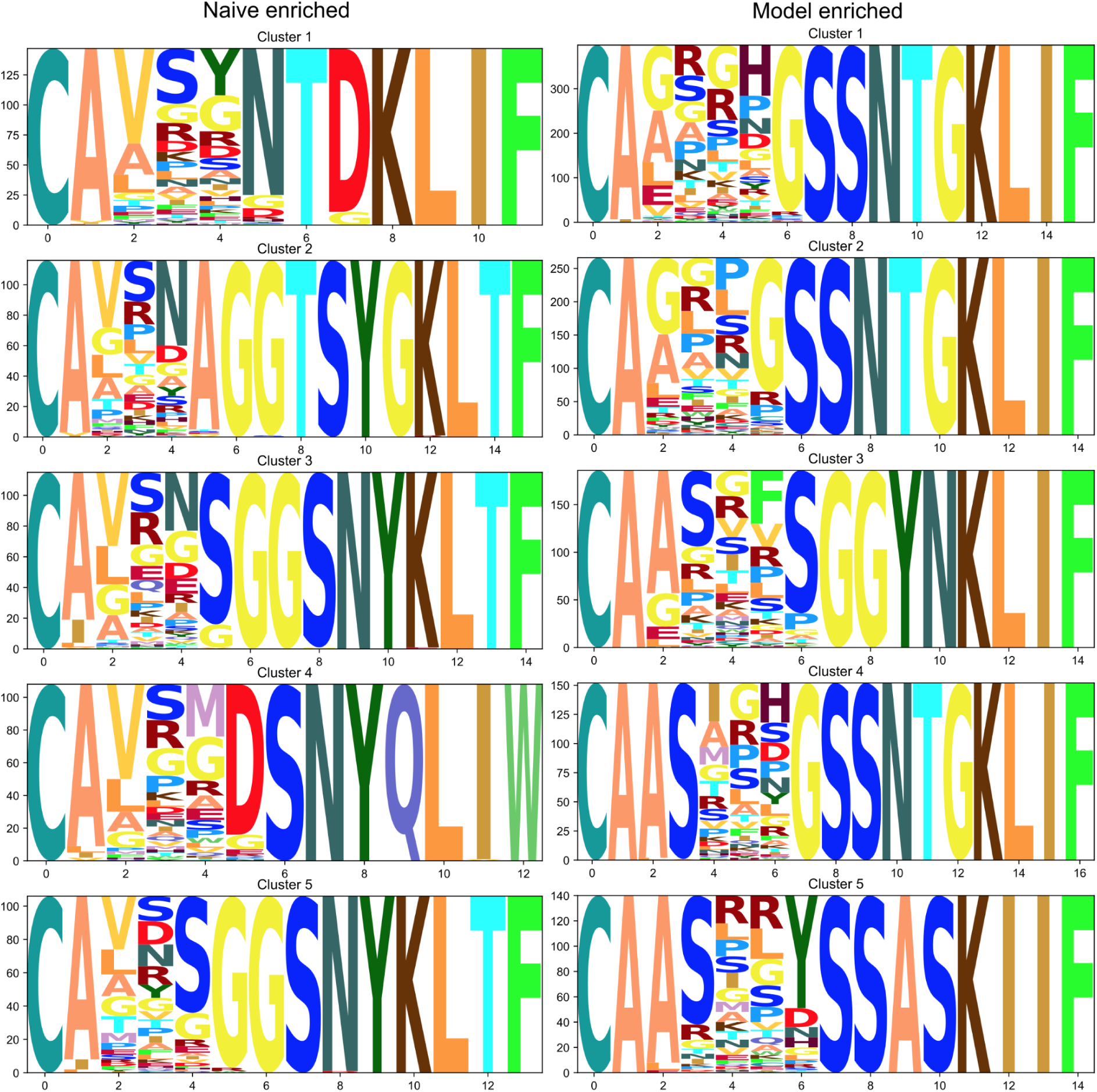
Same as **Supplementary Figure 2** but comparing TCR-CDR3α clusters. In naive T-cells enriched clusters there are a lot of flexible poly-G motifs. In model-enriched clusters there are a lot of SS motifs.

**Supplementary Figure 10.**
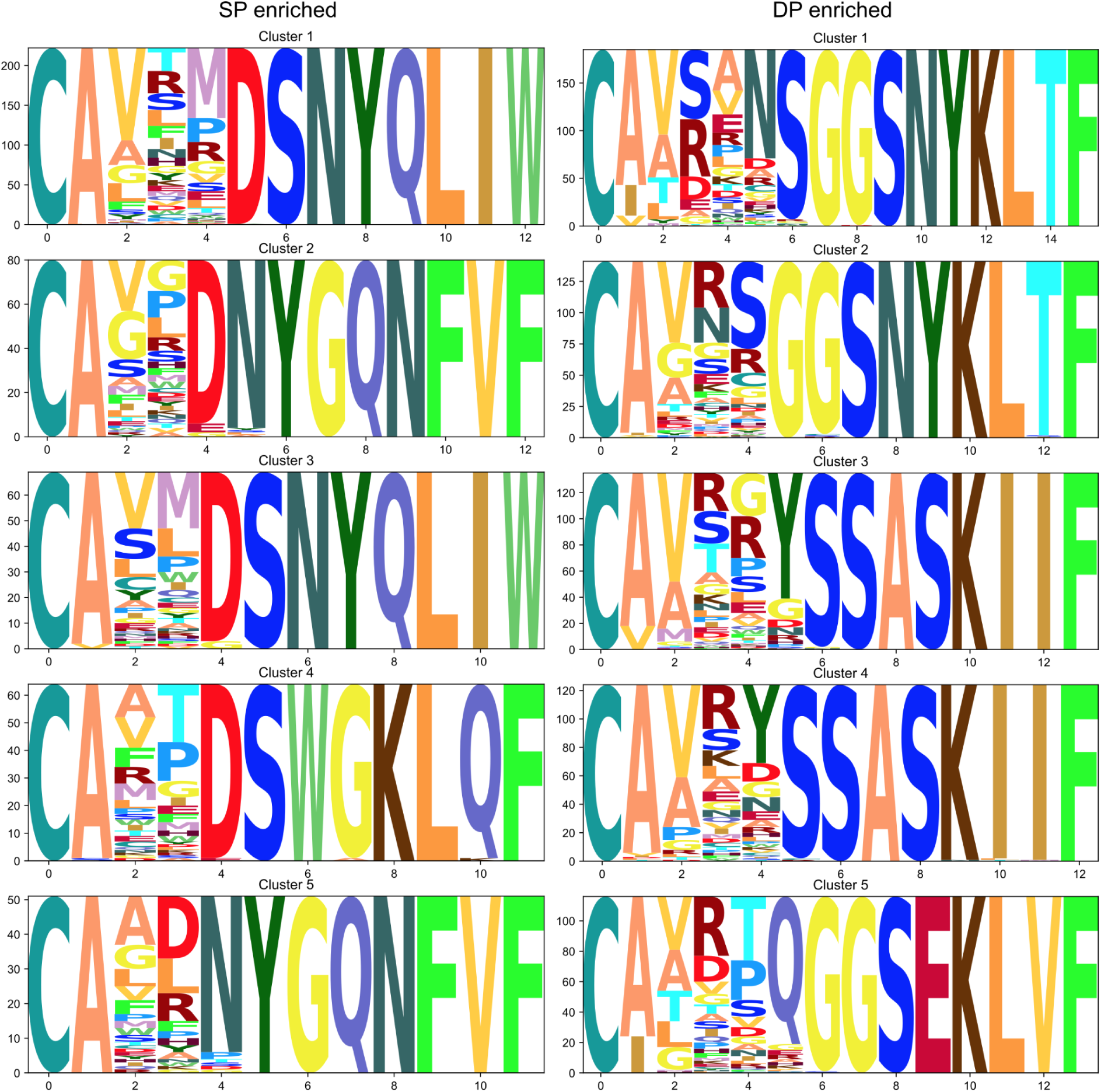
Same as in **Supplementary Figure 2**, but comparing CD8+ single-positive (SP) as a proxy for positively selected TCR-CDR3α motifs and double-positive (DP) as a proxy for pre-selection thymocytes.

**Supplementary Figure 11.**
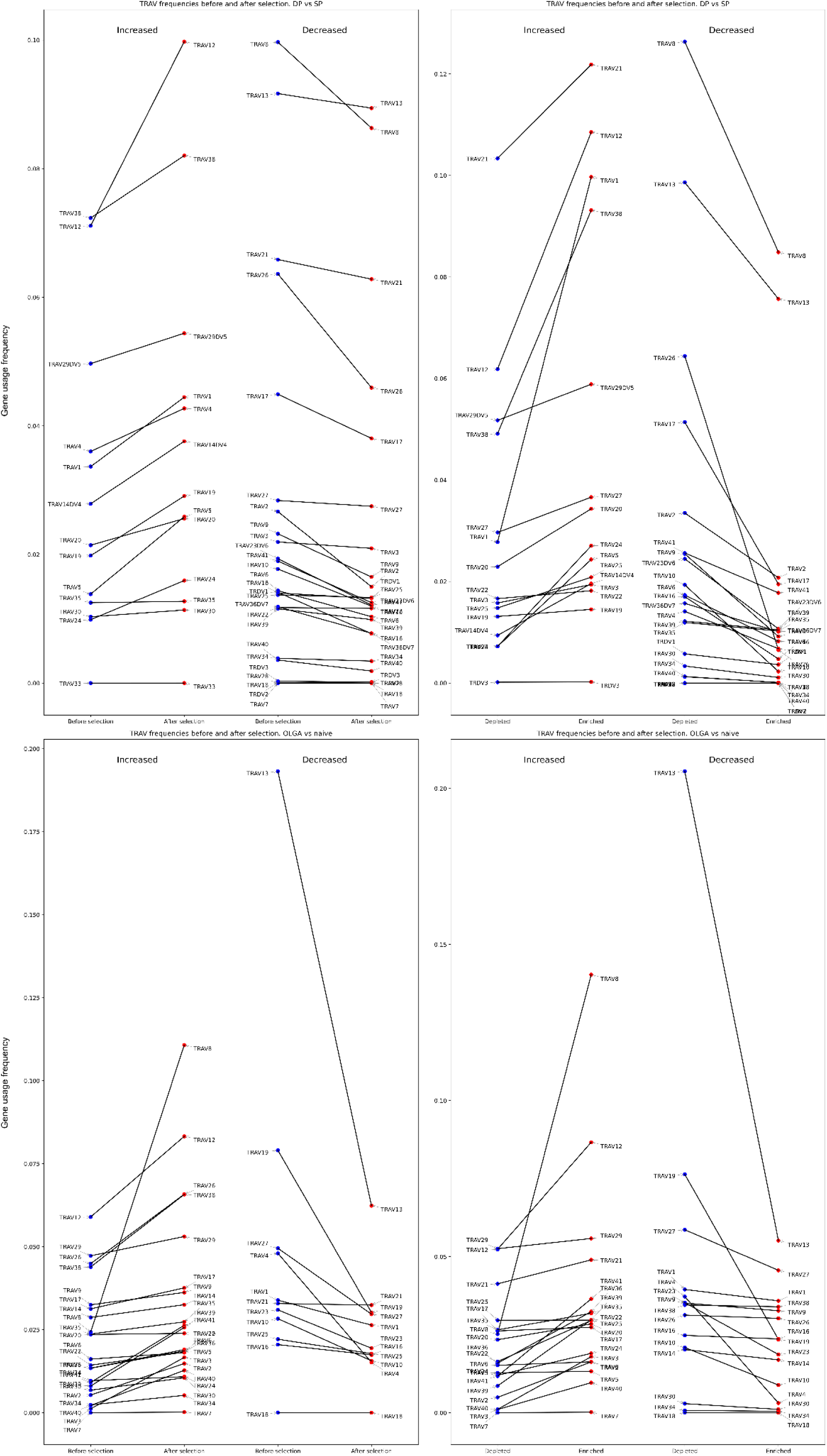
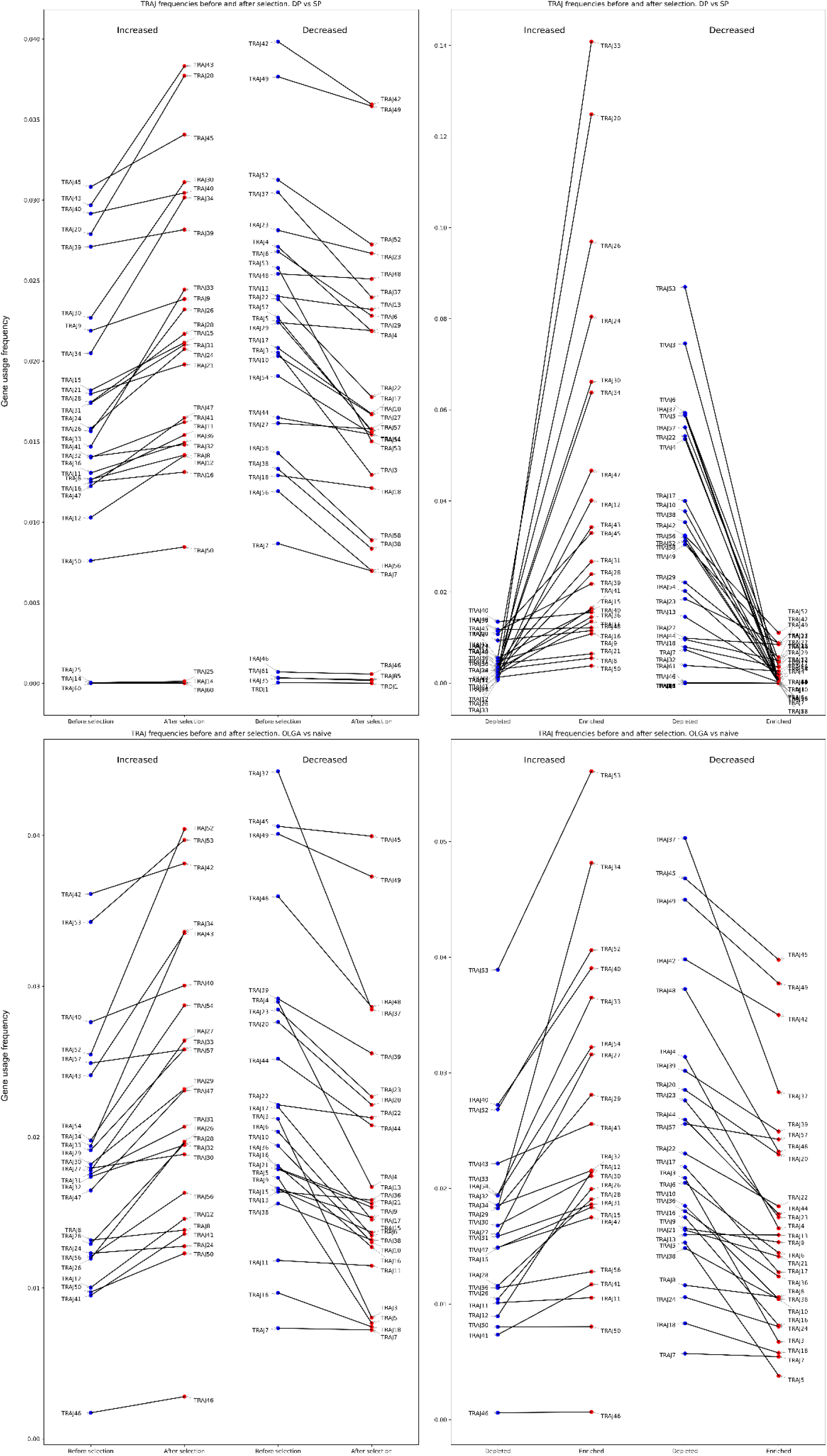
TCRα V and J gene usage before and after selection. Analysis was carried out for entire TCR repertoires and for the enriched and depleted TCRα clusters.

**Supplementary Figure 12.**
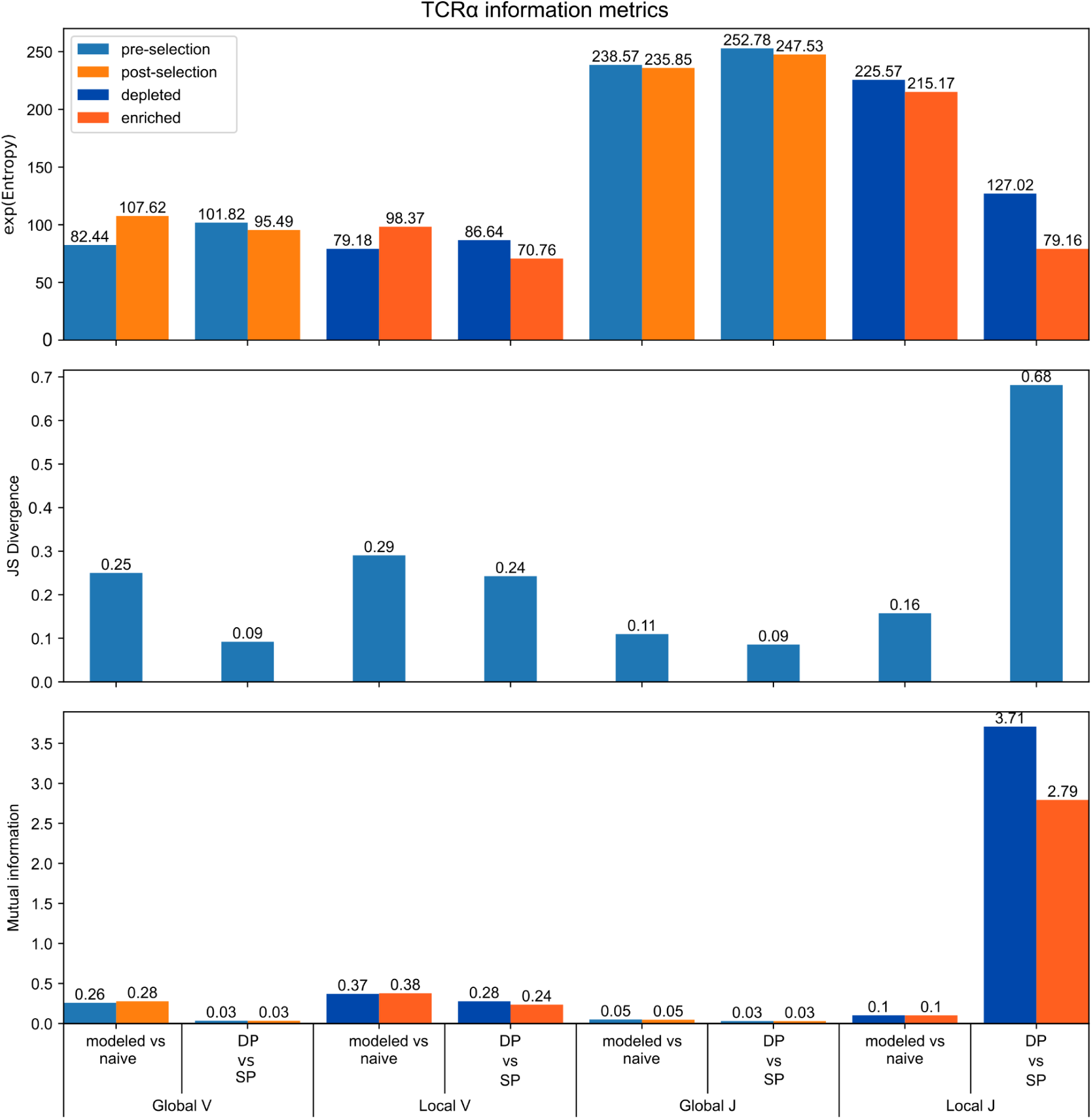
TCRα V and J gene usage information metrics. Analysis was carried out for a whole repertoire (Global) and for enriched pre-/post- selection repertoires (Local).

**Supplementary Table 1.** Metadata of datasets used in the study, number of detected clonotypes (unique VDJ junctions) and number of donors is reported.

| Dataset | Chain | # of clones analyzed | # of subjects | protocol |
| --- | --- | --- | --- | --- |
| Emerson *et al.* | $\beta$ | 1147250 | 10 | Adaptive |
| Heikkilä *et al.* | $\alpha$ | 1582774 | 10 | Adaptive |
| Qi *et al.*, CD4+ | $\beta$ | 1599217 | 9 | Custom |
| Qi *et al.*, CD8+ | $\beta$ | 1346247 | 9 | Custom |
| Quiniou *et al.* | $\beta$ | 1134755 | 14 | 5' RACE, no UMI |
| Quiniou *et al.* | $\alpha$ | 902723 | 14 | 5' RACE, no UMI |
| Pogorelyy *et al.* | $\beta$ | 3678636 | 6, 2 replicas | 5' RACE, with UMI |

**Supplementary Table 2.** HLA typing for 3 pairs of twins from Pogorelyy *et al*. study, class I and II alleles are reported.

|  | S1 and S2 | P1 and P2 | Q1 and Q2 |
| --- | --- | --- | --- |
| A | 02:01:01/03:01:01 | 02:01:01/02:01:01 | 02:01:01/02:01:01 |
| B | 38:01:01/07:02:01 | 51:01:01/07:02:01 | 13:02:01/44:02:01 |
| C | 07:02:01/12:03:01 | 07:02:01/14:02:01 | 05:01:01/06:02:01 |
| DQB1 | 06:03:01/06:02:01 | 03:01:01/04:02:01 | 02:02:01/06:03:01 |
| DRB1 | 13:01:01/15:01:01 | 08:03:02/08:01:03 | 07:01:01/13:01:01 |
| DRB3 | 1:01:02 | - | 1:01:02 |
| DRB4 | - | - | 1:03 |

### Supplementary Note 1

We performed single-cell data analysis focusing on a set of 10,000 CD8+ α and β TCR-CDR3 sequences that are enriched post-selection similar to what is described in the Results section of the main text.

The following additional 10X scRNA-seq datasets have been analyzed:

https://www.10xgenomics.com/resources/datasets/human-pbmc-from-a-healthy-donor-10-k-cells-multi-v-2-2-standard-5-0-0 - “Dataset #1”

https://www.10xgenomics.com/resources/datasets/10k-human-pbmcs-5-v2-0-chromium-x-without-intronic-reads-2-standard - “Dataset #2”

The number of enriched sequences found in Dataset #1 and Dataset #2 was 119 and 108 respectively for CDR3β, 167 and 111 for CDR3α.

The results for both additional Datasets were similar to what is reported in the main text as shown in figures below.

**Supplementary Figure 13.**
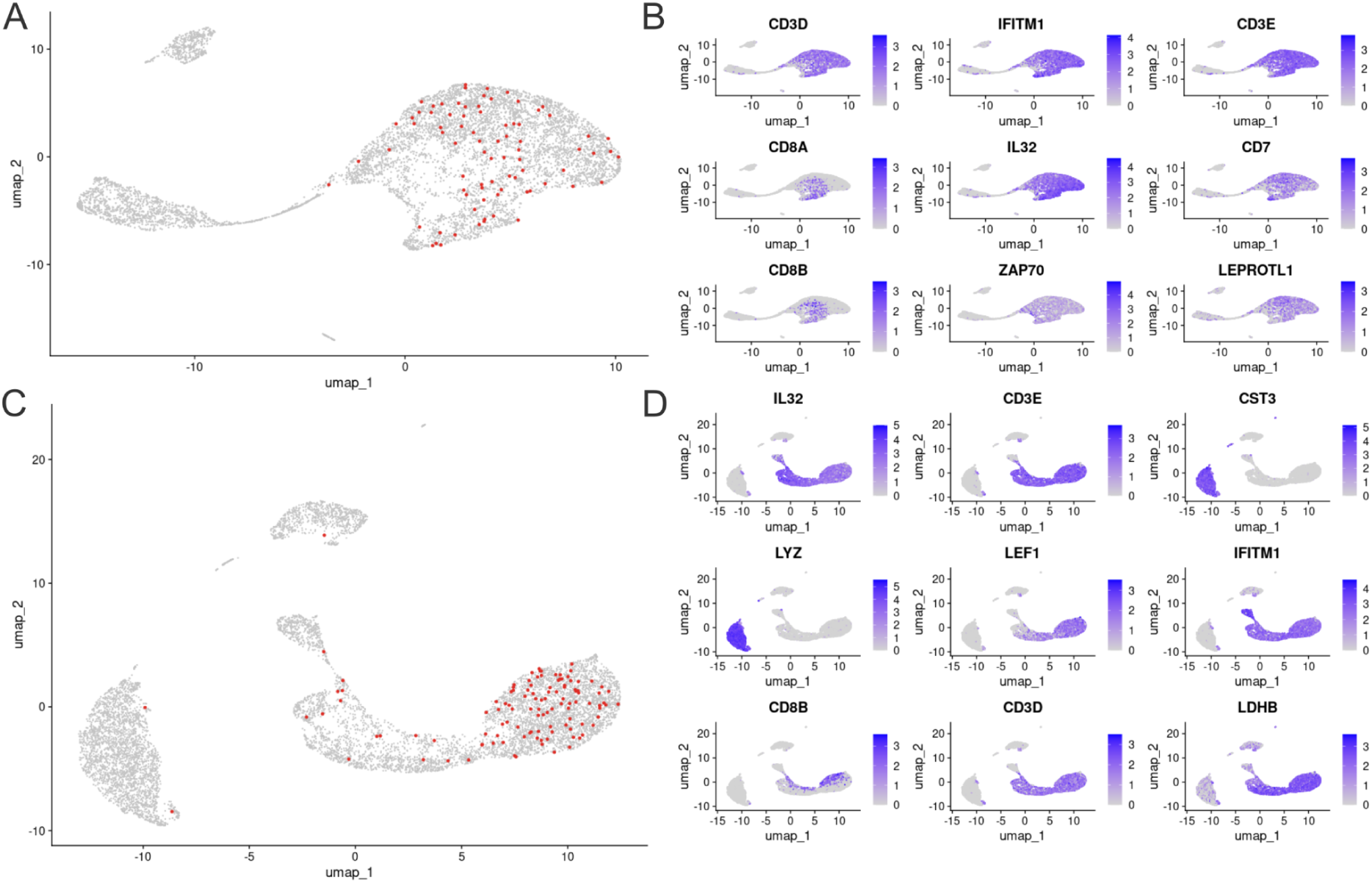
Single cell analysis of enriched CDR3β in 2 additional datasets. **A.** UMAP representation of Dataset #1. **B.** Top 9 differentially expressed genes among cells carrying enriched CDR3β in the Dataset #1. **C.** Enriched CDR3β in UMAP representation of Dataset #2. **D.** Top 9 differentially expressed genes, either enriched or depleted, among cells carrying enriched CDR3β in the Dataset #2.

As expected, with CDR3s enriched in CD8 thymocytes repertoire demonstrated differential expression of CD8+ T-cells markers CD8A and CD8B, highlighting the fact that inferred motifs can predict T-cell phenotype (Supplementary Figure 13B) (Duan and Mukherjee, 2016). Additional markers, such as IFITM1 (CD225) that is enriched are subject to further exploration.

**Supplementary Figure 14.**
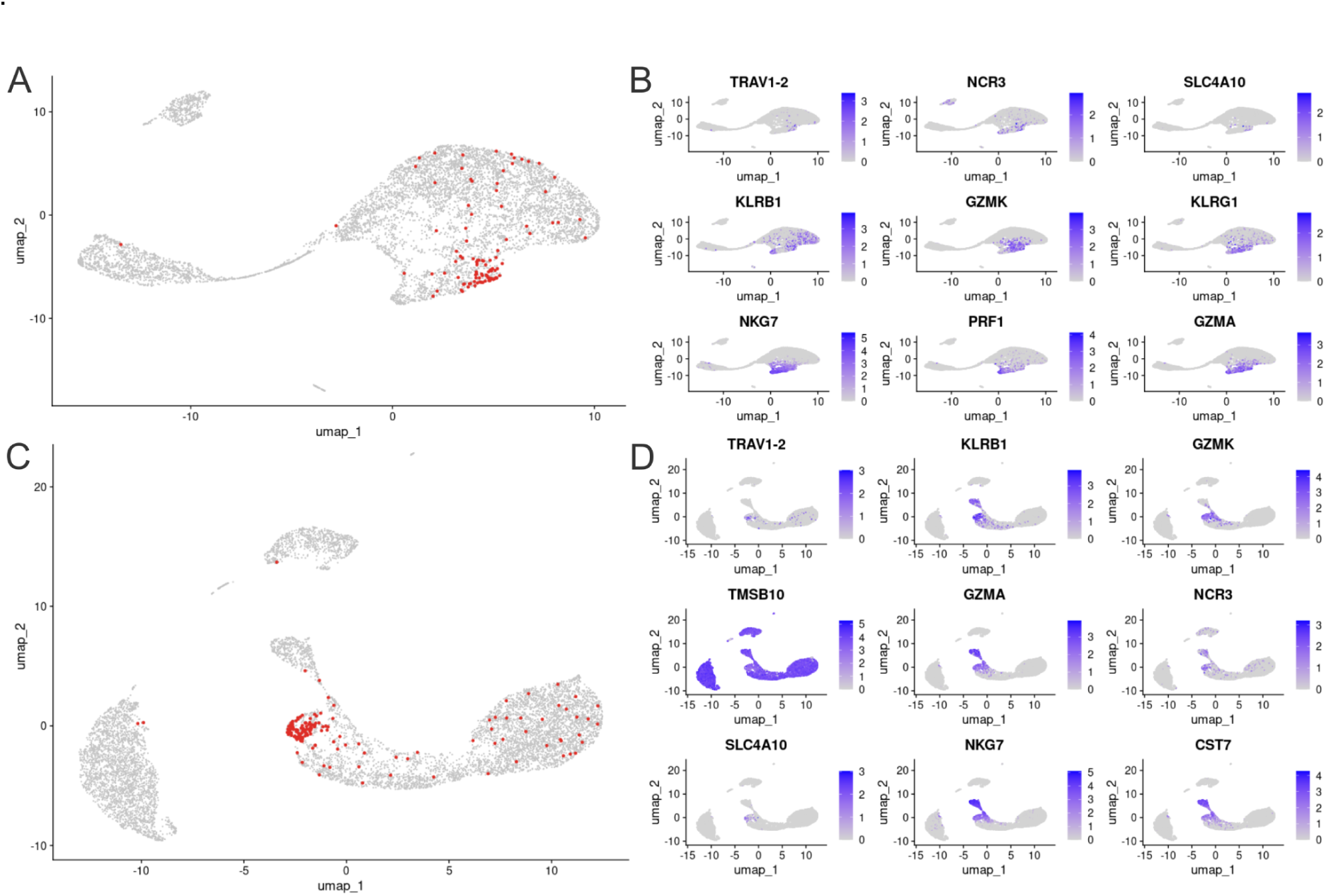
Single-cell analysis of enriched CDR3α in 2 additional datasets. **A.** CDR3α enriched repertoires mapped to single-cell UMAP plot of Dataset #1. **B.** Top 9 differentially expressed genes among cells carrying enriched CDR3α in the Dataset #1. **C.** CDR3α enriched repertoires mapped to single-cell UMAP plot of Dataset #2. **D.** Top 9 differentially expressed genes among cells carrying enriched CDR3α in the Dataset #2.

Results for the analysis of CDR3α were identical to those obtained in the main part of the article (Figure 2). Top 9 most differentially expressed genes in this group represented the MAIT cells signature (Godfrey et al., 2019). Similar to the Result section, cells carrying the enriched CDR3α were densely located in the compartmentalised part of the UMAP. Overall, these findings support the robustness of lineage commitment guided by the TCR sequence motifs.

### Supplementary Note 2

We reproduced the results of “HLA allele affects the selection” section using two additional datasets of twin TCR repertoires reported in (Zvyagin et al., 2014)(dataset#2) and (Kasatskaya et al., 2020) (dataset#3).

The number of TCR clonotypes (unique VDJ junctions) in samples from each twin varied from 22k to 171k. We calculated Jensen–Shannon divergence for them in the same manner as for the dataset reported in the main text. This time the clustering for relative twins was much less pronounced. It is especially notable for sample a1, from which the number of clonotypes was smallest in the dataset (Supplementary Figure 15, Supplementary table 3). Thus, we speculate that lack of clustering may be due to insufficient sample size of data being analyzed.

**Supplementary Figure 15.**
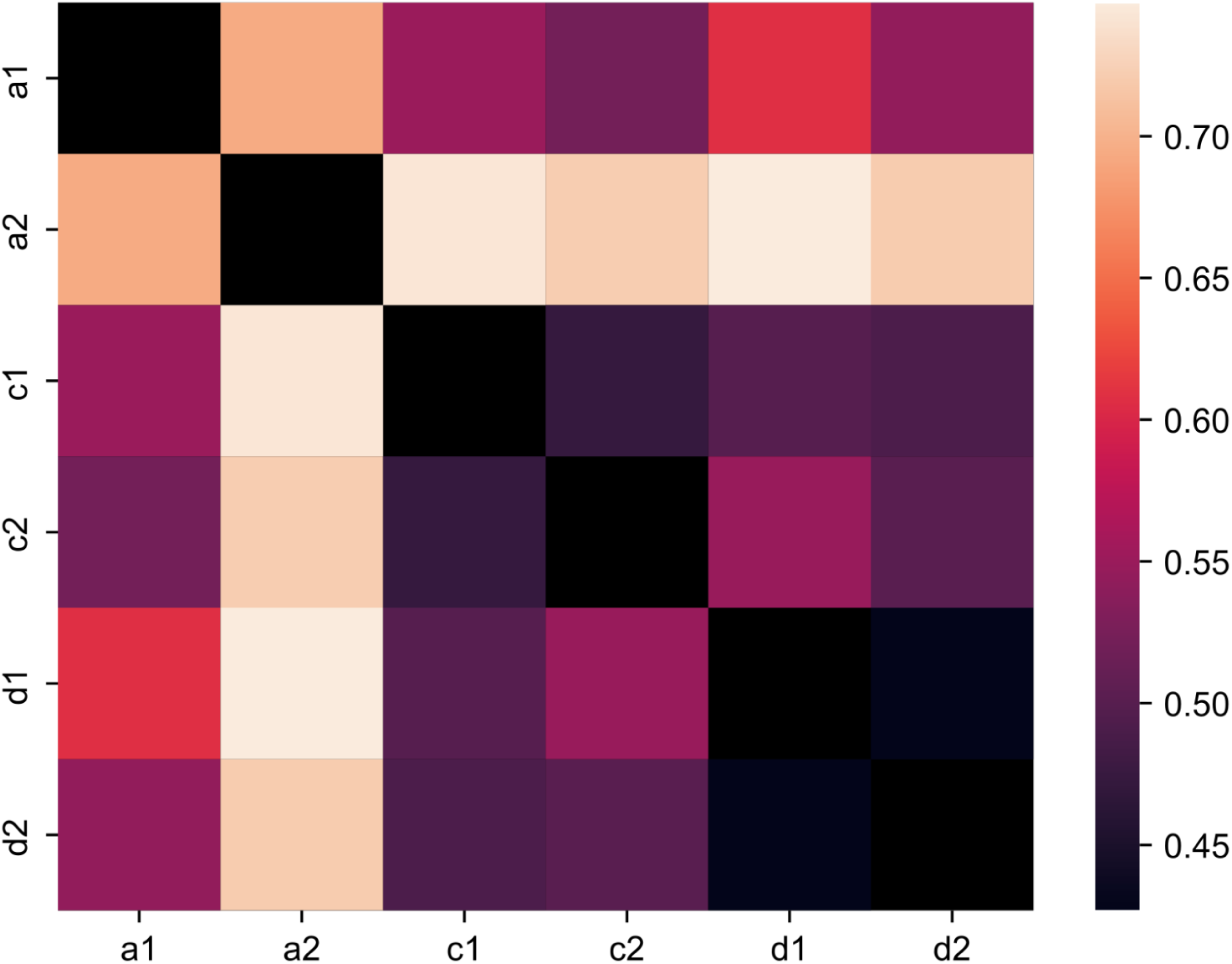
Jensen-Shannon divergence for frequencies of positively-selected (enriched against a background sample produced using VDJ rearrangement model) CDR3β clusters in additional twin dataset (dataset #2).

**Supplementary Table 3.** Sample sizes of the additional twin dataset (dataset#2).

| Sample | TCRα clonotypes | TCRβ clonotypes |
| --- | --- | --- |
| a1 | 39378 | 53997 |
| a2 | 19134 | 22430 |
| c1 | 129837 | 171502 |
| c2 | 69964 | 78934 |
| d1 | 71951 | 91228 |
| d2 | 75272 | 166666 |

We then decided to check whether or not sample size affects clustering and what sample size is required for clustering to appear. To do so, we took the initial twin’s dataset (analyzed in a Result section “Twins CDR3 clusterization”) and then gradually reduced its sample size. Five replicas of sample size reduction were conducted. Then the mean Jensen-Shannon divergence was calculated for related twins and for unrelated twins along with confidence intervals.

**Supplementary Figure 16.**
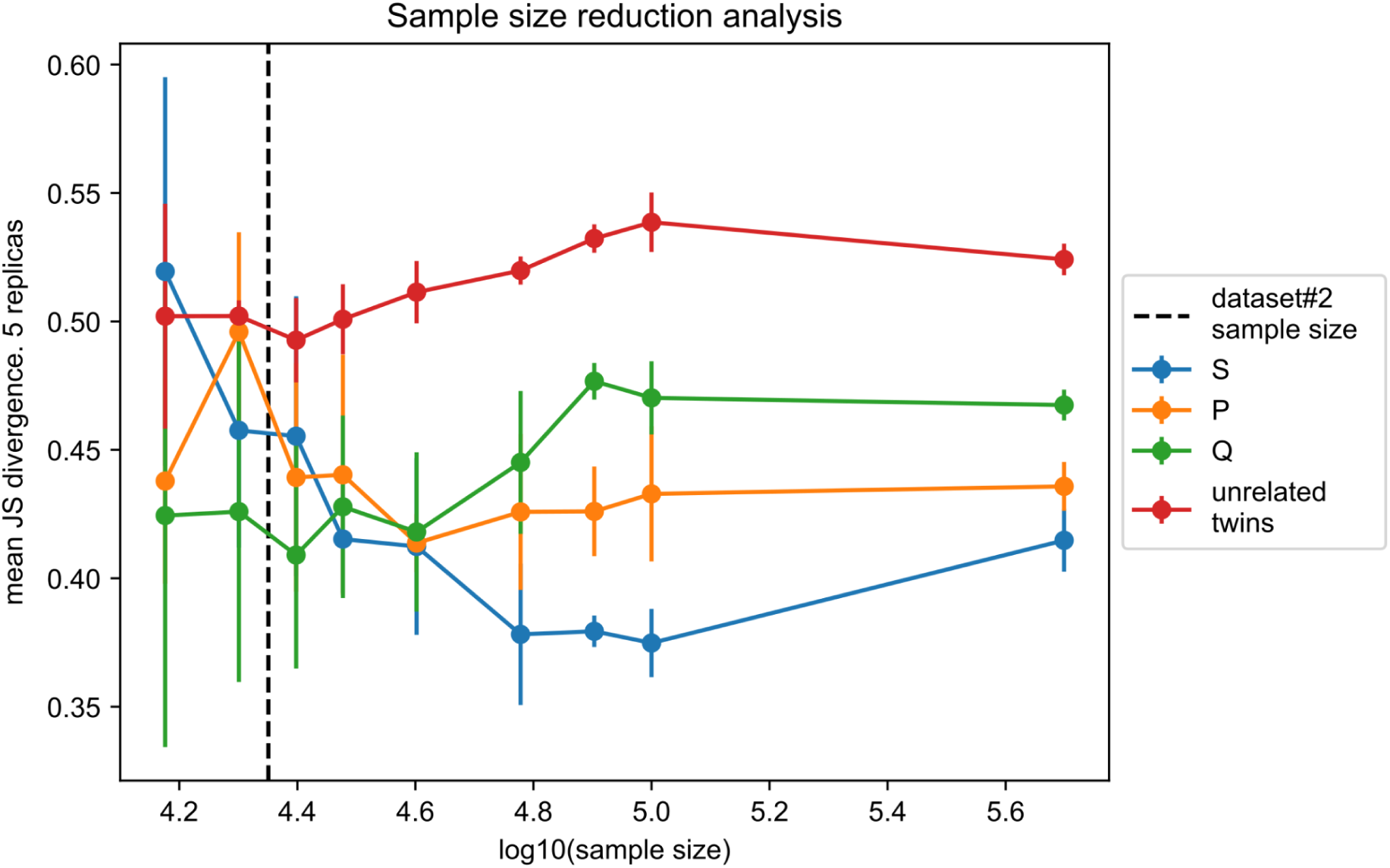
Sample size affects the ability to distinguish relative and non-relative twins. Each point represents Jensen-Shannon divergence between relative twin pairs S, P, and Q or between unrelated twins averaged for five sample size reduction replicas. Error bars show confidence intervals calculated for 5 sample size reduction replicas. Dashed line represents the sample size of datasets#2.

Apparently, for successful HLA allele effect detection, at least 40k TCR clones from each sample are needed (Supplementary Figure 16). From this sample size, confidence intervals of relative twins no longer intersect with confidence intervals of unrelated twins. The sample size of datasets#2 (22k) is smaller than this value; thus, we can suggest that a modest sample size is the main reason why we failed to detect relative twins clustering in dataset#2.

Additionally, we ran clustering for motif frequences in dataset#3, which has CD4+ CDR3β from two pairs of twins. This dataset spans three subsets of T-cells for each sample: T-regulatory cells (Treg), recent thymic emigrants (RTEs) and all other cells. However, since the sample size of each subset was too small to detect clustering, we combined them together. (**Supplementary Table 4**).

We indeed detected the same effect as in the first dataset (Supplementary figure 17). However the effect on model enriched (depleted) TCR was not detected (Supplementary figure 18). It is still an open question whether this is due to not-sufficient sample size or due to absence of HLA allele effect on TCRs which did not pass through the selection in case of CD4+ cells.

**Supplementary Table 4.** Sample sizes of twins dataset#3 in terms of unique TCR beta sequences.

|  | Treg | RTE | N_nonRTE | Total |
| --- | --- | --- | --- | --- |
| J_1 | 27556 | 16765 | 72080 | 116401 |
| J_2 | 46922 | 38389 | 44957 | 130268 |
| C_1 | 33779 | 56979 | 38385 | 129143 |
| C_2 | 24927 | 64670 | 56000 | 145597 |

**Supplementary Figure 17.**
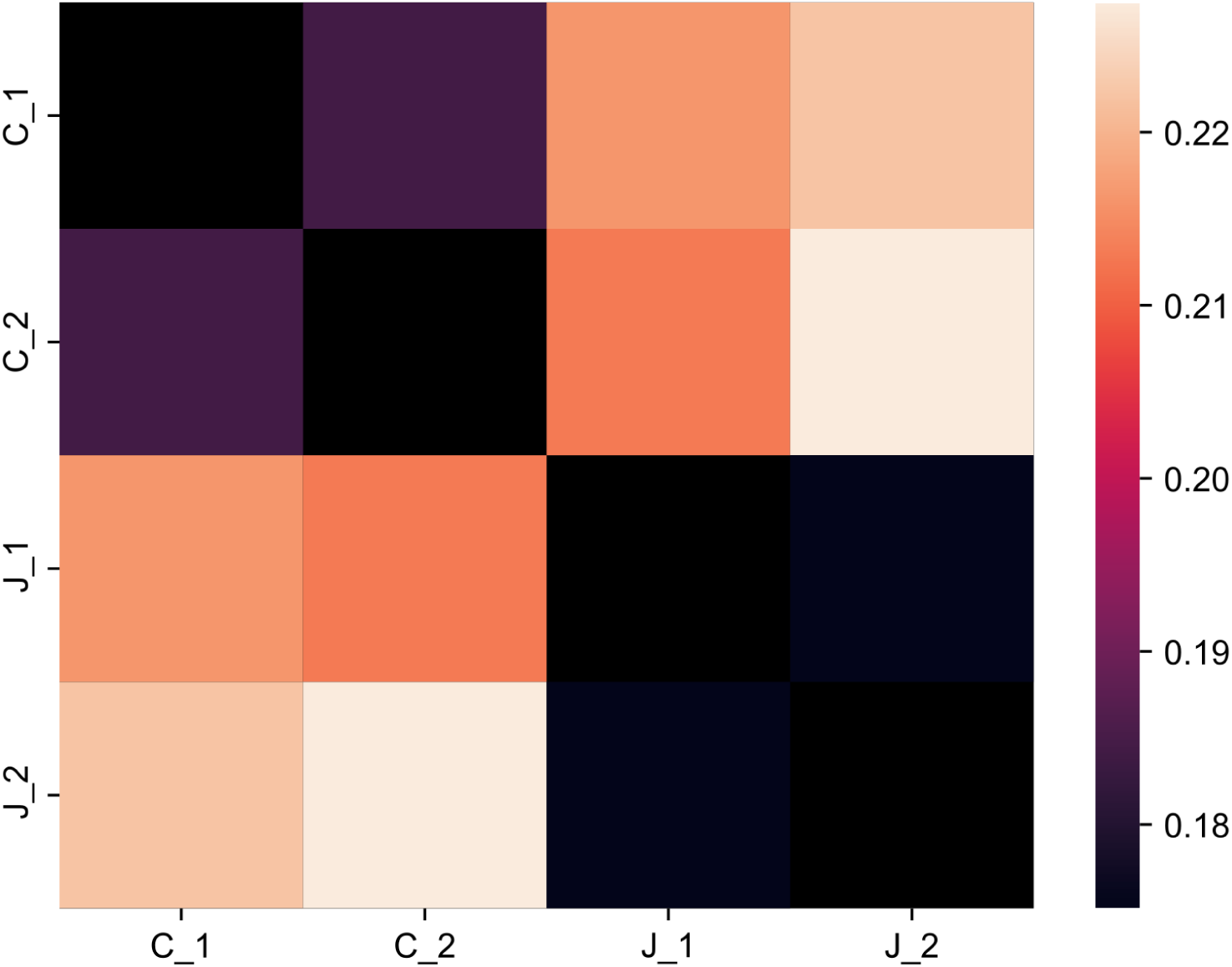
Jensen-Shannon divergence for CDR3β clusters in dataset#3 enriched (positively selected) against a background dataset of TCRs generated using VDJ rearrangement model.

**Supplementary Figure 18.**
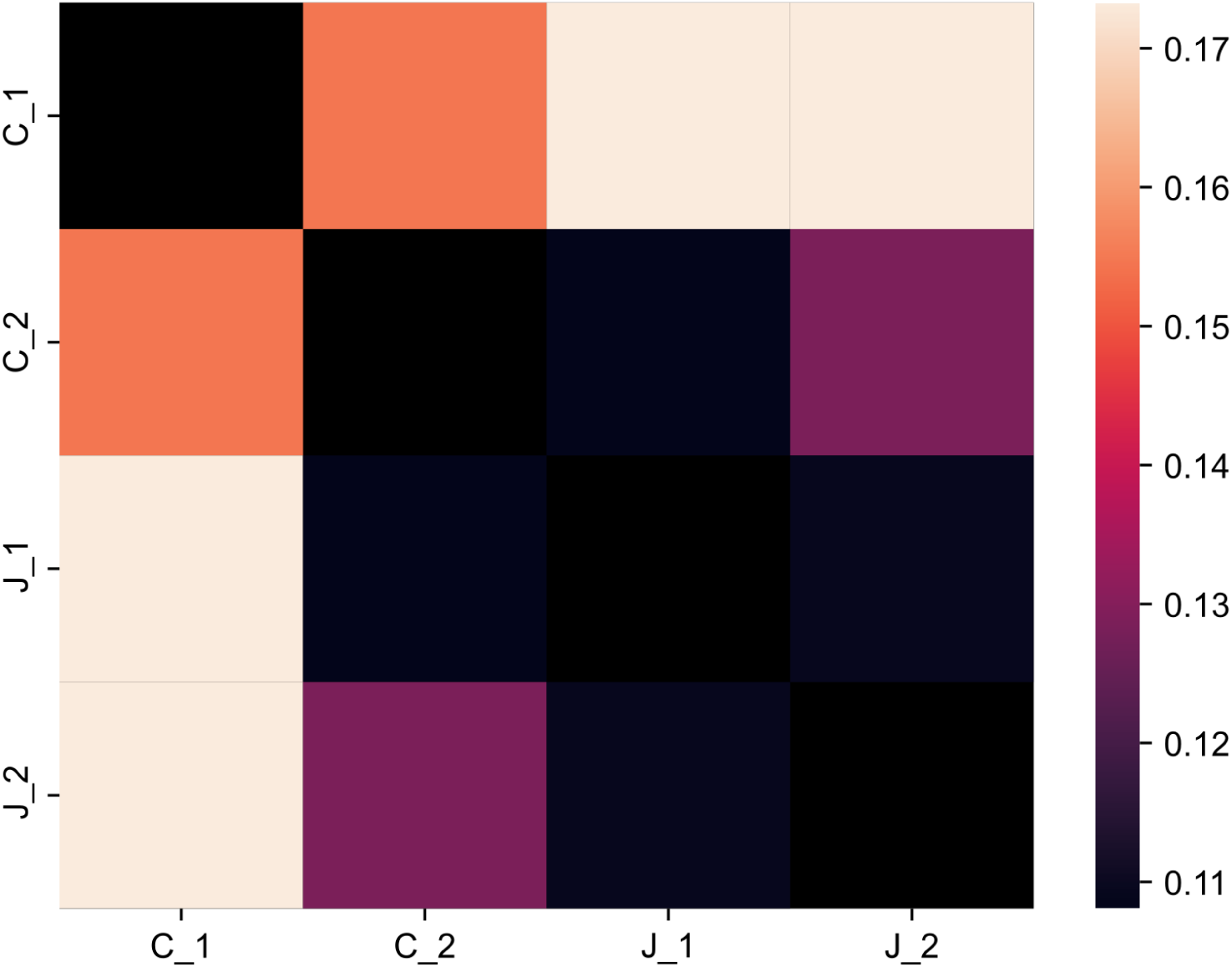
Jensen-Shannon divergence for CDR3β clusters in dataset#3 depleted (negatively selected) against a background dataset of TCRs generated using VDJ rearrangement model.

## References

Ashby, K.M., Hogquist, K.A., 2023. A guide to thymic selection of T cells. Nat. Rev. Immunol. 1–15. 10.1038/s41577-023-00911-8

Barennes, P., Quiniou, V., Shugay, M., Egorov, E.S., Davydov, A.N., Chudakov, D.M., Uddin, I., Ismail, M., Oakes, T., Chain, B., Eugster, A., Kashofer, K., Rainer, P.P., Darko, S., Ransier, A., Douek, D.C., Klatzmann, D., Mariotti-Ferrandiz, E., 2021. Benchmarking of T cell receptor repertoire profiling methods reveals large systematic biases. Nat. Biotechnol. 39, 236–245. 10.1038/s41587-020-0656-3

Benichou, J., Ben-Hamo, R., Louzoun, Y., Efroni, S., 2012. Rep-Seq: uncovering the immunological repertoire through next-generation sequencing. Immunology 135, 183–191. 10.1111/j.1365-2567.2011.03527.x

Bolotin, D.A., Mamedov, I.Z., Britanova, O.V., Zvyagin, I.V., Shagin, D., Ustyugova, S.V., Turchaninova, M.A., Lukyanov, S., Lebedev, Y.B., Chudakov, D.M., 2012. Next generation sequencing for TCR repertoire profiling: Platform-specific features and correction algorithms. Eur. J. Immunol. 42, 3073–3083. 10.1002/eji.201242517

Bolze, A., Neveux, I., Schiabor Barrett, K.M., White, S., Isaksson, M., Dabe, S., Lee, W., Grzymski, J.J., Washington, N.L., Cirulli, E.T., 2022. HLA-A∗03:01 is associated with increased risk of fever, chills, and stronger side effects from Pfizer-BioNTech COVID-19 vaccination. Hum. Genet. Genomics Adv. 3, 100084. 10.1016/j.xhgg.2021.100084

Bosselut, R., 2019. T cell antigen recognition: Evolution-driven affinities. Proc. Natl. Acad. Sci. U. S. A. 116, 21969–21971. 10.1073/pnas.1916129116

Britanova, O.V., Putintseva, E.V., Shugay, M., Merzlyak, E.M., Turchaninova, M.A., Staroverov, D.B., Bolotin, D.A., Lukyanov, S., Bogdanova, E.A., Mamedov, I.Z., Lebedev, Y.B., Chudakov, D.M., 2014. Age-Related Decrease in TCR Repertoire Diversity Measured with Deep and Normalized Sequence Profiling. J. Immunol. 192, 2689–2698. 10.4049/jimmunol.1302064

Brown, A.J., White, J., Shaw, L., Gross, J., Slabodkin, A., Kushner, E., Greiff, V., Matsuda, J., Gapin, L., Scott-Browne, J., Kappler, J., Marrack, P., 2024. MHC heterozygosity limits T cell receptor variability in CD4 T cells. Sci. Immunol. 9, eado5295. 10.1126/sciimmunol.ado5295

Bykov, S., Asher, S., 2010. Raman studies of Solution Polyglycine Conformations. J. Phys. Chem. B 114, 6636–6641. 10.1021/jp100082n

Camaglia, F., Ryvkin, A., Greenstein, E., Reich-Zeliger, S., Chain, B., Mora, T., Walczak, A.M., Friedman, N., 2023. Quantifying changes in the T cell receptor repertoire during thymic development. eLife 12, e81622. 10.7554/eLife.81622

Chowell, D., Morris, L.G.T., Grigg, C.M., Weber, J.K., Samstein, R.M., Makarov, V., Kuo, F., Kendall, S.M., Requena, D., Riaz, N., Greenbaum, B., Carroll, J., Garon, E., Hyman, D.M., Zehir, A., Solit, D., Berger, M., Zhou, R., Rizvi, N.A., Chan, T.A., 2018. Patient HLA class I genotype influences cancer response to checkpoint blockade immunotherapy. Science 359, 582–587. 10.1126/science.aao4572

Cole, D., 2013. Re-Directing CD4+ T Cell Responses with the Flanking Residues of MHC Class II-Bound Peptides: The Core is Not Enough. Front. Immunol. 4.

De Simone, G., Mazza, E.M.C., Cassotta, A., Davydov, A.N., Kuka, M., Zanon, V., De Paoli, F., Scamardella, E., Metsger, M., Roberto, A., Pilipow, K., Colombo, F.S., Tenedini, E., Tagliafico, E., Gattinoni, L., Mavilio, D., Peano, C., Price, D.A., Singh, S.P., Farber, J.M., Serra, V., Cucca, F., Ferrari, F., Orrù, V., Fiorillo, E., Iannacone, M., Chudakov, D.M., Sallusto, F., Lugli, E., 2019. CXCR3 Identifies Human Naive CD8+ T Cells with Enhanced Effector Differentiation Potential. J. Immunol. Baltim. Md 1950 203, 3179–3189. 10.4049/jimmunol.1901072

Duan, L., Mukherjee, E., 2016. Janeway’s Immunobiology, Ninth Edition. Yale J. Biol. Med. 89, 424–425.

Dupic, T., Marcou, Q., Walczak, A.M., Mora, T., 2019. Genesis of the αβ T-cell receptor. PLOS Comput. Biol. 15, e1006874. 10.1371/journal.pcbi.1006874

Elhanati, Y., Murugan, A., Callan, C.G., Mora, T., Walczak, A.M., 2014. Quantifying selection in immune receptor repertoires. Proc. Natl. Acad. Sci. 111, 9875–9880. 10.1073/pnas.1409572111

Farber, D.L., Yudanin, N.A., Restifo, N.P., 2014. Human memory T cells: generation, compartmentalization and homeostasis. Nat. Rev. Immunol. 14, 24–35. 10.1038/nri3567

Feng, Y., Van Der Veeken, J., Shugay, M., Putintseva, E.V., Osmanbeyoglu, H.U., Dikiy, S., Hoyos, B.E., Moltedo, B., Hemmers, S., Treuting, P., Leslie, C.S., Chudakov, D.M., Rudensky, A.Y., 2015. A mechanism for expansion of regulatory T-cell repertoire and its role in self-tolerance. Nature 528, 132–136. 10.1038/nature16141

Godfrey, D.I., Koay, H.-F., McCluskey, J., Gherardin, N.A., 2019. The biology and functional importance of MAIT cells. Nat. Immunol. 20, 1110–1128. 10.1038/s41590-019-0444-8

H, T., Tm, G., Jr, M., W, C., Y, T., Re, D., D, P., Sj, C., Wh, H., Cl, D., L, T., Cm, W., G, G., Jj, G., 2020. Determinants governing T cell receptor α/β-chain pairing in repertoire formation of identical twins. PubMed.

Heikkilä, N., Sormunen, S., Mattila, J., Härkönen, T., Knip, M., Ihantola, E.-L., Kinnunen, T., Mattila, I.P., Saramäki, J., Arstila, T.P., 2021. Generation of self-reactive, shared T-cell receptor α chains in the human thymus. J. Autoimmun. 119, 102616. 10.1016/j.jaut.2021.102616

Huang, Y.-N., Patel, N.A., Mehta, J.H., Ginjala, S., Brodin, P., Gray, C.M., Patel, Y.M., Cowell, L.G., Burkhardt, A.M., Mangul, S., 2022. Data Availability of Open T-Cell Receptor Repertoire Data, a Systematic Assessment. Front. Syst. Biol. 2, 918792. 10.3389/fsysb.2022.918792

Isacchini, G., Quiniou, V., Barennes, P., Mhanna, V., Vantomme, H., Stys, P., Mariotti-Ferrandiz, E., Klatzmann, D., Walczak, A.M., Mora, T., Nourmohammad, A., 2024. Local and Global Variability in Developing Human T-Cell Repertoires. PRX Life 2, 013011. 10.1103/PRXLife.2.013011

Isacchini, G., Walczak, A.M., Mora, T., Nourmohammad, A., 2021. Deep generative selection models of T and B cell receptor repertoires with soNNia. Proc. Natl. Acad. Sci. 118, e2023141118. 10.1073/pnas.2023141118

Ishigaki, K., Lagattuta, K.A., Luo, Y., James, E.A., Buckner, J.H., Raychaudhuri, S., 2022. HLA autoimmune risk alleles restrict the hypervariable region of T cell receptors. Nat. Genet. 54, 393–402. 10.1038/s41588-022-01032-z

Kalos, M., June, C.H., 2013. Adoptive T cell Transfer for Cancer Immunotherapy in the Era of Synthetic Biology. Immunity 39, 10.1016/j.immuni.2013.07.002. 10.1016/j.immuni.2013.07.002

Karnaukhov, V.K., Shcherbinin, D.S., Chugunov, A.O., Chudakov, D.M., Efremov, R.G., Zvyagin, I.V., Shugay, M., 2024. Structure-based prediction of T cell receptor recognition of unseen epitopes using TCRen. Nat. Comput. Sci. 4, 510–521. 10.1038/s43588-024-00653-0

Kasatskaya, S.A., Ladell, K., Egorov, E.S., Miners, K.L., Davydov, A.N., Metsger, M., Staroverov, D.B., Matveyshina, E.K., Shagina, I.A., Mamedov, I.Z., Izraelson, M., Shelyakin, P.V., Britanova, O.V., Price, D.A., Chudakov, D.M., n.d. Functionally specialized human CD4+ T-cell subsets express physicochemically distinct TCRs. eLife 9, e57063. 10.7554/eLife.57063

Kidera, A., Konishi, Y., Oka, M., Ooi, T., Scheraga, H.A., 1985. Statistical analysis of the physical properties of the 20 naturally occurring amino acids. J. Protein Chem. 4, 23–55. 10.1007/BF01025492

Kosmrlj, A., Jha, A.K., Huseby, E.S., Kardar, M., Chakraborty, A.K., 2008. How the thymus designs antigen-specific and self-tolerant T cell receptor sequences. Proc. Natl. Acad. Sci. U. S. A. 105, 16671–16676. 10.1073/pnas.0808081105

Leich, E., Maier, C., Bomben, R., Vit, F., Bosi, A., Horn, H., Gattei, V., Ott, G., Rosenwald, A., Zamò, A., 2021. Follicular lymphoma subgroups with and without t(14;18) differ in their N-glycosylation pattern and IGHV usage. Blood Adv. 5, 4890–4900. 10.1182/bloodadvances.2021005081

Li, X., Mizsei, R., Tan, K., Mallis, R.J., Duke-Cohan, J.S., Akitsu, A., Tetteh, P.W., Dubey, A., Hwang, W., Wagner, G., Lang, M.J., Arthanari, H., Wang, J., Reinherz, E.L., 2021. Pre–T cell receptors topologically sample self-ligands during thymocyte β-selection. Science 371, 181–185. 10.1126/science.abe0918

Lindeboom, R.G.H., Worlock, K.B., Dratva, L.M., Yoshida, M., Scobie, D., Wagstaffe, H.R., Richardson, L., Wilbrey-Clark, A., Barnes, J.L., Kretschmer, L., Polanski, K., Allen-Hyttinen, J., Mehta, P., Sumanaweera, D., Boccacino, J.M., Sungnak, W., Elmentaite, R., Huang, N., Mamanova, L., Kapuge, R., Bolt, L., Prigmore, E., Killingley, B., Kalinova, M., Mayer, M., Boyers, A., Mann, A., Swadling, L., Woodall, M.N.J., Ellis, S., Smith, C.M., Teixeira, V.H., Janes, S.M., Chambers, R.C., Haniffa, M., Catchpole, A., Heyderman, R., Noursadeghi, M., Chain, B., Mayer, A., Meyer, K.B., Chiu, C., Nikolić, M.Z., Teichmann, S.A., 2024. Human SARS-CoV-2 challenge uncovers local and systemic response dynamics. Nature 631, 189–198. 10.1038/s41586-024-07575-x

Logunova, N.N., Kriukova, V.V., Shelyakin, P.V., Egorov, E.S., Pereverzeva, A., Bozhanova, N.G., Shugay, M., Shcherbinin, D.S., Pogorelyy, M.V., Merzlyak, E.M., Zubov, V.N., Meiler, J., Chudakov, D.M., Apt, A.S., Britanova, O.V., 2020. MHC-II alleles shape the CDR3 repertoires of conventional and regulatory naïve CD4+ T cells. Proc. Natl. Acad. Sci. 117, 13659–13669. 10.1073/pnas.2003170117

Lu, J., Van Laethem, F., Bhattacharya, A., Craveiro, M., Saba, I., Chu, J., Love, N.C., Tikhonova, A., Radaev, S., Sun, X., Ko, A., Arnon, T., Shifrut, E., Friedman, N., Weng, N.-P., Singer, A., Sun, P.D., 2019. Molecular constraints on CDR3 for thymic selection of MHC-restricted TCRs from a random pre-selection repertoire. Nat. Commun. 10, 1019. 10.1038/s41467-019-08906-7

Markov, P.V., Pybus, O.G., 2015. Evolution and Diversity of the Human Leukocyte Antigen(HLA). Evol. Med. Public Health 2015, 1. 10.1093/emph/eou033

Marshall, R.D., 1972. Glycoproteins. Annu. Rev. Biochem. 41, 673–702. 10.1146/annurev.bi.41.070172.003325

Nakonechnaya, T.O., Moltedo, B., Putintseva, E.V., Leyn, S., Bolotin, D.A., Britanova, O.V., Shugay, M., Chudakov, D.M., 2024. Convergence, plasticity, and tissue residence of regulatory T cell response via TCR repertoire prism. eLife 12, RP89382. 10.7554/eLife.89382

Noble, J.A., Valdes, A.M., 2011. Genetics of the HLA Region in the Prediction of Type 1 Diabetes. Curr. Diab. Rep. 11, 533–542. 10.1007/s11892-011-0223-x

Pogorelyy, M.V., Minervina, A.A., Touzel, M.P., Sycheva, A.L., Komech, E.A., Kovalenko, E.I., Karganova, G.G., Egorov, E.S., Komkov, A.Y., Chudakov, D.M., Mamedov, I.Z., Mora, T., Walczak, A.M., Lebedev, Y.B., 2018. Precise tracking of vaccine-responding T cell clones reveals convergent and personalized response in identical twins. Proc. Natl. Acad. Sci. 115, 12704–12709. 10.1073/pnas.1809642115

Pogorelyy, M.V., Shugay, M., 2019. A Framework for Annotation of Antigen Specificities in High-Throughput T-Cell Repertoire Sequencing Studies. Front. Immunol. 10, 2159. 10.3389/fimmu.2019.02159

Pospelova, M., Safonova, Y., 2022. Analyzing patterns of tyrosine sulfation in naive antibody repertoires. 10.1101/2022.12.13.520330

Qi, Q., Liu, Y., Cheng, Y., Glanville, J., Zhang, D., Lee, J.-Y., Olshen, R.A., Weyand, C.M., Boyd, S.D., Goronzy, J.J., 2014. Diversity and clonal selection in the human T-cell repertoire. Proc. Natl. Acad. Sci. 111, 13139–13144. 10.1073/pnas.1409155111

Quiniou, V., Barennes, P., Mhanna, V., Stys, P., Vantomme, H., Zhou, Z., Martina, F., Coatnoan, N., Barbie, M., Pham, H.-P., Clémenceau, B., Vie, H., Shugay, M., Six, A., Brandao, B., Mallone, R., Mariotti-Ferrandiz, E., Klatzmann, D., 2023. Human thymopoiesis produces polyspecific CD8+ α/β T cells responding to multiple viral antigens. eLife 12, e81274. 10.7554/eLife.81274

Robert, P.A., Kunze-Schumacher, H., Greiff, V., Krueger, A., 2021. Modeling the Dynamics of T-Cell Development in the Thymus. Entropy 23, 437. 10.3390/e23040437

Rosati, E., Dowds, C.M., Liaskou, E., Henriksen, E.K.K., Karlsen, T.H., Franke, A., 2017. Overview of methodologies for T-cell receptor repertoire analysis. BMC Biotechnol. 17, 61. 10.1186/s12896-017-0379-9

Rothenberg, E.V., 2023. The β-selection step shapes T-cell identity. Nature 613, 440–442. 10.1038/d41586-023-00025-0

Rozemuller, E., Eckle, S.B.G., McLaughlin, I., Penning, M., Mulder, W., de Bruin, H., van Wageningen, S., 2021. MR1 encompasses at least six allele groups with coding region alterations. HLA 98, 509–516. 10.1111/tan.14390

Rubelt, F., Busse, C.E., Bukhari, S.A.C., Bürckert, J.-P., Mariotti-Ferrandiz, E., Cowell, L.G., Watson, C.T., Marthandan, N., Faison, W.J., Hershberg, U., Laserson, U., Corrie, B.D., Davis, M.M., Peters, B., Lefranc, M.-P., Scott, J.K., Breden, F., Luning Prak, E.T., Kleinstein, S.H., 2017. Adaptive Immune Receptor Repertoire Community recommendations for sharing immune-repertoire sequencing data. Nat. Immunol. 18, 1274–1278. 10.1038/ni.3873

Schneider, D., Veelken, H., Jumaa, H., 2012. The Functional Role of Acquired N-Linked Glycosylation Sites On Follicular Lymphoma B Cell Antigen Receptors. Blood 120, 2704. 10.1182/blood.V120.21.2704.2704

Schumacher, T.N., Scheper, W., Kvistborg, P., 2019. Cancer Neoantigens. Annu. Rev. Immunol. 37, 173–200. 10.1146/annurev-immunol-042617-053402

Sethna, Z., Elhanati, Y., Callan, C.G., Walczak, A.M., Mora, T., 2019. OLGA: fast computation of generation probabilities of B- and T-cell receptor amino acid sequences and motifs. Bioinformatics 35, 2974–2981. 10.1093/bioinformatics/btz035

Shugay, M., Bagaev, D.V., Turchaninova, M.A., Bolotin, D.A., Britanova, O.V., Putintseva, E.V., Pogorelyy, M.V., Nazarov, V.I., Zvyagin, I.V., Kirgizova, V.I., Kirgizov, K.I., Skorobogatova, E.V., Chudakov, D.M., 2015. VDJtools: Unifying Post-analysis of T Cell Receptor Repertoires. PLOS Comput. Biol. 11, e1004503. 10.1371/journal.pcbi.1004503

Stadinski, B.D., Shekhar, K., Gómez-Touriño, I., Jung, J., Sasaki, K., Sewell, A.K., Peakman, M., Chakraborty, A.K., Huseby, E.S., 2016. Hydrophobic CDR3 residues promote the development of self-reactive T cells. Nat. Immunol. 17, 946–955. 10.1038/ni.3491

Sun, S., Hu, Y., Ao, M., Shah, P., Chen, J., Yang, W., Jia, X., Tian, Y., Thomas, S., Zhang, H., 2019. N-GlycositeAtlas: a database resource for mass spectrometry-based human N-linked glycoprotein and glycosylation site mapping. Clin. Proteomics 16, 35. 10.1186/s12014-019-9254-0

TCRen: predicting TCR recognition of unseen epitopes based on residue-level pairwise statistical potential | bioRxiv [WWW Document], n.d. URL https://www.biorxiv.org/content/10.1101/2022.02.15.480516v2.abstract (accessed 1.22.24).

Treiner, E., Duban, L., Bahram, S., Radosavljevic, M., Wanner, V., Tilloy, F., Affaticati, P., Gilfillan, S., Lantz, O., 2003. Selection of evolutionarily conserved mucosal-associated invariant T cells by MR1. Nature 422, 164–169. 10.1038/nature01433

Yates, A., 2014. Theories and Quantification of Thymic Selection. Front. Immunol. 5.

Yin, R., Ribeiro-Filho, H.V., Lin, V., Gowthaman, R., Cheung, M., Pierce, B.G., 2023. TCRmodel2: high-resolution modeling of T cell receptor recognition using deep learning. Nucleic Acids Res. 51, W569–W576. 10.1093/nar/gkad356

Zhu, D., McCarthy, H., Ottensmeier, C.H., Johnson, P., Hamblin, T.J., Stevenson, F.K., 2002. Acquisition of potential N-glycosylation sites in the immunoglobulin variable region by somatic mutation is a distinctive feature of follicular lymphoma. Blood 99, 2562–2568. 10.1182/blood.v99.7.2562

Zvyagin, I.V., Pogorelyy, M.V., Ivanova, M.E., Komech, E.A., Shugay, M., Bolotin, D.A., Shelenkov, A.A., Kurnosov, A.A., Staroverov, D.B., Chudakov, D.M., Lebedev, Y.B., Mamedov, I.Z., 2014. Distinctive properties of identical twins’ TCR repertoires revealed by high-throughput sequencing. Proc. Natl. Acad. Sci. 111, 5980–5985. 10.1073/pnas.1319389111

